# Distinct Cdk9-phosphatase switches act at the beginning and end of elongation by RNA polymerase II

**DOI:** 10.1101/2020.06.14.150847

**Authors:** Pabitra K. Parua, Sampada Kalan, Bradley Benjamin, Miriam Sansó, Robert P. Fisher

**Affiliations:** Department of Oncological Sciences, Icahn School of Medicine at Mount Sinai, New York, NY, 10029-6574, USA

## Abstract

Reversible phosphorylation of Pol II and accessory factors helps order the transcription cycle. Here we define two kinase-phosphatase switches that operate at different points in human transcription. Cdk9/cyclin T1 (P-TEFb) catalyzes inhibitory phosphorylation of PP1 and PP4 complexes that localize to 3’ and 5’ ends of genes, respectively, and have overlapping but distinct specificities for Cdk9-dependent phosphorylations of Spt5, a factor instrumental in promoter-proximal pausing and elongation-rate control. PP1 dephosphorylates an Spt5 carboxy-terminal repeat (CTR), but not Spt5-Ser666, a site between KOW motifs 4 and 5, whereas PP4 can target both sites. In vivo, Spt5-CTR phosphorylation decreases as transcription complexes pass the cleavage and polyadenylation signal (CPS) and increases upon PP1 depletion, consistent with a PP1 function in termination first uncovered in yeast. Depletion of PP4-complex subunits increases phosphorylation of both Ser666 and the CTR, and promotes redistribution of promoter-proximally paused Pol II into gene bodies. These results suggest that switches comprising Cdk9 and either PP4 or PP1 govern pause release and the elongation-termination transition, respectively.

---

The transcription cycle of RNA polymerase II (Pol II) is divided into discrete phases of initiation, elongation and termination. This process is regulated by cyclin-dependent kinases (CDKs) that generate stage-specific patterns of phosphorylation on the carboxy-terminal domain (CTD) of the Pol II large subunit Rpb1 ^1,2^. Differential phosphorylation of the CTD, which consists of heptad repeats of consensus sequence Y_1_S_2_P_3_T_4_S_5_P_6_S_7_, inscribes a “CTD code” ^3–5^ that is read by factors and enzymes that preferentially bind the modified CTD, in part to coordinate RNA-processing and chromatin modification with transcription ^6^. Concurrently, CDKs phosphorylate many other targets to control progression through the transcription cycle ^7–9^.

A promoter-proximal pause soon after the transition from initiation to elongation is a rate-limiting step in transcription of many Pol II-dependent genes in metazoans ^10,11^. This pause is established within the first ~100 nucleotides (nt) downstream of the transcription start site (TSS) by recruitment of the DRB-sensitivity inducing factor (DSIF)—a heterodimer of Spt4 and Spt5 subunits conserved in all eukaryotes—and a metazoan-specific negative elongation factor (NELF) ^12^. In human cells, DSIF and NELF recruitment (and thus, pause establishment) depends on activity of Cdk7, a component of transcription initiation factor TFIIH ^13–16^, whereas pause release depends on the Cdk9/cyclin T1 complex, also known as positive transcription elongation factor b (P-TEFb) ^17^, which phosphorylates residues in Pol II, Spt5, NELF and other components of the paused complex, to convert it into an active elongation complex from which NELF is displaced ^18,19^. P-TEFb and its orthologs in yeast are the major CDKs active during the elongation phase of Pol II transcription, phosphorylating Spt5 to enable its function as a processivity factor ^20^, and stimulating elongation rate by 3-4-fold ^21,22^.

Pol II undergoes a second, 3’ pause downstream of the cleavage and polyadenylation signal (CPS) ^23^; this slowing manifests as a peak of Pol II occupancy in chromatin immunoprecipitation (ChIP) or run-on transcription profiles. Pol II paused downstream of the CPS becomes heavily phosphorylated on Ser2 of the CTD, which occurs as a consequence—rather than a cause—of its decreased elongation rate ^24,25^. Pausing and Ser2 phosphorylation (pSer2) in turn promote recruitment of factors needed for mRNA 3’-end maturation and termination ^26^. We recently uncovered a regulatory circuit in fission yeast comprising Cdk9, the protein phosphatase 1 (PP1) isoform Dis2, and their common enzymatic target, Spt5, with the potential to switch Pol II from rapid elongation to a paused state permissive for termination ^27^. During processive elongation, Cdk9 phosphorylates the Spt5 CTD and keeps Dis2 inactive by phosphorylating its carboxy-terminal region. As elongation complexes traverse the CPS, the Spt5 CTD is dephosphorylated dependent on activity of Dis2, which is a subunit of the cleavage and polyadenylation factor (CPF) ^28^; the drop in phospho-Spt5 precedes an increase in pSer2 over the 3’ pause site. Inactivation of Cdk9 or Dis2 leads to opposite effects downstream of the CPS—more rapid termination, or more extensive read-through transcription indicating a termination defect, respectively ^21,27^. The effect of Cdk9 inhibition was recapitulated by an *spt5* mutation that prevented Spt5-CTD phosphorylation ^29^. Recently, human PP1 and its regulatory subunit PNUTS were implicated in Spt5-CTR dephosphorylation and Pol II deceleration downstream of the CPS ^30,31^, suggesting conservation of this mechanism.

Here we show first that the entire Cdk9-PP1-Spt5 switch is conserved in human cells. Two PP1 catalytic-subunit isoforms and two residues of Spt5 were among targets of human P-TEFb we identified in a chemical-genetic screen ^9^. Cdk9 inhibition diminishes phosphorylation of PP1γ on a known inhibitory site, and of Spt5 on carboxy-terminal repeat region 1 (CTR1), whereas depletion of PP1 increases steady-state levels of CTR1 phosphorylation (pCTR1). In unperturbed cells, pCTR1 drops, Pol II accumulates and pSer2 increases downstream of the CPS—the same relationships seen in fission yeast ^27^. The Cdk9 substrate screen also identified Spt5-Ser666, a site outside the CTRs in a region linking Kyrpides-Ouzounis-Woese (KOW) motifs 4 and 5 ^9^. Although Ser666 phosphorylation (pSer666) depends on Cdk9, it is resistant to dephosphorylation by PP1, and pSer666 and pCTR1 are distributed differently on chromatin: pSer666 increases beyond the promoter-proximal pause and is retained downstream of the CPS. We identify a second site of Cdk9-mediated inhibitory phosphorylation in PP4R2, a regulatory subunit of the protein phosphatase 4 (PP4) complex. PP4 can dephosphorylate pSer666 in vitro, in contrast to PP1, and is excluded from chromatin near the 3’-ends of genes where PP1γ occupancy is maximal, potentially explaining why pSer666 is not removed downstream of the CPS. PP4 depletion increases pSer666 and pCTR1 levels and attenuates promoter-proximal pausing in vivo. Therefore, Cdk9 phosphorylates multiple sites on Spt5 while restraining activity of two phosphatases with different site-specificities and chromatin distributions, to generate diverse spatial patterns of Spt5 phosphorylation and possibly to support discrete functions at different steps of the transcription cycle.

## Results

### A conserved kinase-phosphatase switch in transcription

In fission yeast, Cdk9 phosphorylates the Spt5 CTD ^32^ and the inhibitory Thr316 residue of PP1 isoform Dis2 ^27^. As Pol II traverses the CPS, Spt5-CTD phosphorylation decreases dependent on Dis2 activity, and pSer2-containing Pol II accumulates with Spt5 in a 3’-paused complex poised for termination ^27,29^. We asked if this switch is conserved in human cells, where two PP1 catalytic-subunit isoforms were identified in a chemical-genetic screen for direct Cdk9 substrates ^9^. We validated PP1γ-Thr311 as a Cdk9-dependent phosphorylation site by two approaches. First, we treated green fluorescent protein (GFP)-tagged PP1γ, expressed in HCT116 cells and immobilized with anti-GFP antibodies, with purified Cdk9/cyclin T1, followed by immunoblotting with an antibody specific for PP1 isoforms phosphorylated on their carboxy-terminal inhibitory sites. Increased signal after Cdk9 treatment of wild-type PP1γ but not PP1γ^T311A^ suggests that P-TEFb can indeed phosphorylate this residue in vitro (Fig. 1a).

**Fig. 1.**
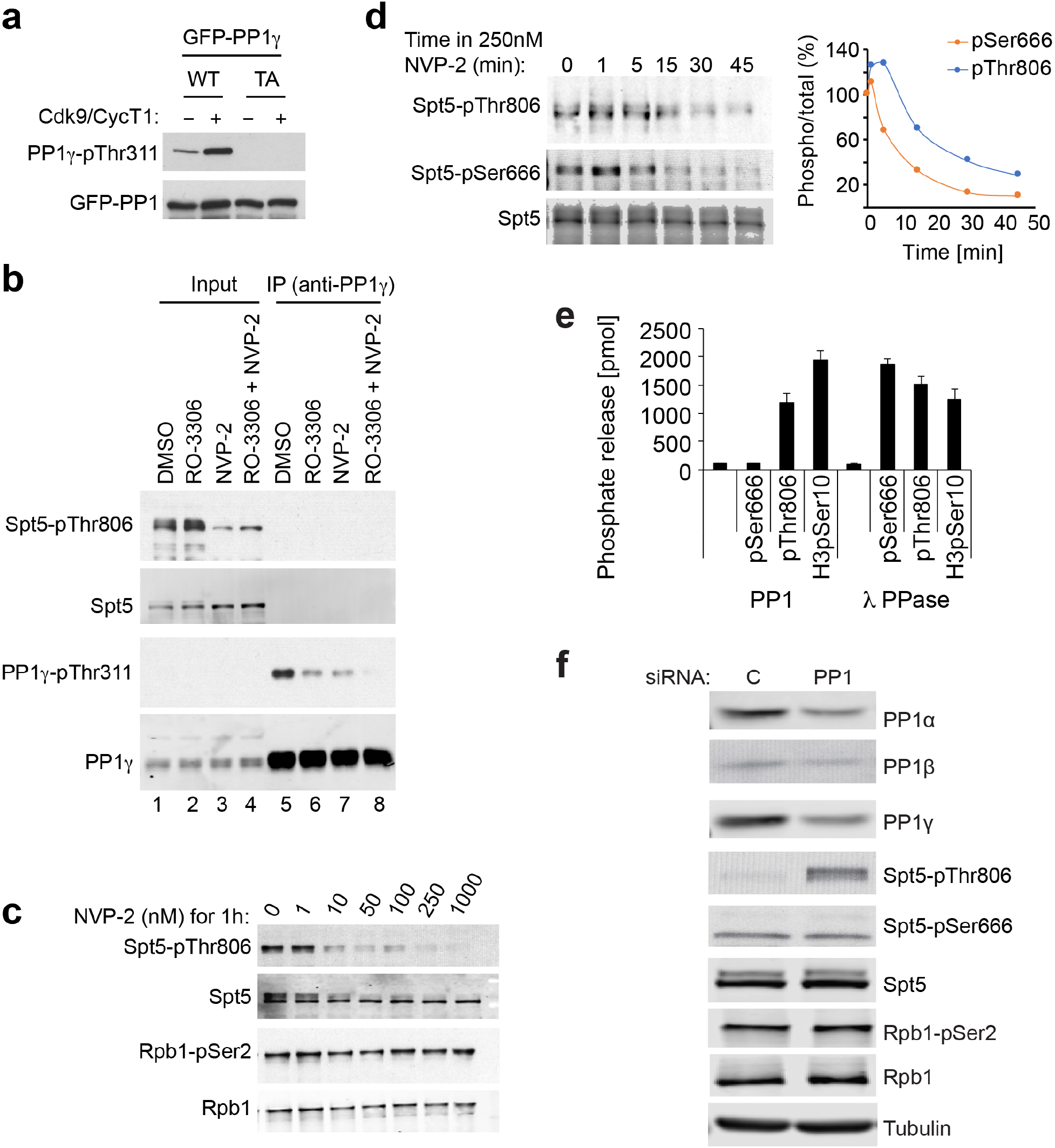
A Cdk9-PP1 switch governing Spt5 phosphorylation is conserved in human cells. **a** Purified, recombinant Cdk9/cyclin T1 phosphorylates wild-type (WT) GFP-PP1γ, expressed in human cells and recovered by anti-GFP immunoprecipitation, but not a Thr3111→Ala (TA) mutant variant. Phosphorylation was detected with antibody specific for the carboxy-terminal phosphorylation site (Thr320) in PP1α isoform, analogous to Thr311 of PP1γ. **b** Inhibition of Cdk9 or Cdk1 diminishes PP1γ-inhibitory phosphorylation in human cells. HCT116 cells were treated with DMSO, a Cdk1 inhibitor (RO-3306), a Cdk9 inhibitor (NVP-2), or both, as indicated. Extracts were analyzed by direct immunoblotting (lanes 1-4), or anti-PP1γ immunoprecipitation followed by immunoblotting (lanes 5-8), with the indicated antibodies. **c** Cdk9 inhibition diminishes phosphorylation of Spt5-Thr806 but not Ser2 of the Pol II CTD. HCT116 cells were treated with the indicated concentrations of NVP-2 for 1 hr and extracts were immunoblotted with the indicated antibodies. **d** HCT116 cells were treated with 250 nM NVP-2 for indicated times and extracts were immunoblotted with antibodies specific for Spt5, Spt5-pThr806 and Spt5-pSer666, as indicated. Immunoblot signals were quantified with ImageJ software. **e** Spt5-derived phosphopeptides containing pSer666 or pThr806 or a control histone H3-derived phosphopeptide containing pSer10, as indicated, were incubated with purified PP1 or lambda phosphatase, as indicated, and phosphate release was measured colorimetrically. Error bars indicate + standard deviation from mean (s.d.) of three biological replicates. **f** HCT116 cells were transfected with an siRNA cocktail targeting all three PP1 catalytic-subunit isoforms or a control (scrambled) siRNA, as indicated, and extracts were immunoblotted for the indicated proteins or protein modifications.

Next we asked if phosphorylation of this site depends on Cdk9 in vivo. Treatment of HCT116 cells with the Cdk9-selective inhibitor NVP-2 ^33^ diminished reactivity of immunoprecipitated PP1γ with the phospho-PP1 antibody to about the same extent as did a Cdk1-selective inhibitor (RO-3306), whereas combined treatment with NVP-2 and RO-3306 nearly abolished the signal (Fig. 1b), suggesting roughly equal contributions of the two CDKs to negative regulation of PP1γ in vivo. We surmise that PP1γ, like fission yeast Dis2 ^27,34^, is a regulatory component shared between the cell-division and transcription machineries.

CTR1 of human Spt5 contains multiple repeats of consensus sequence G-S-Q/R-T-P, including Thr806—a Cdk9 target site detected in our screen ^9^—and is analogous to the CTD of the fission yeast protein (Supplementary Fig. 1a). After a 1-hr treatment with 10-50 nM NVP-2, pThr806 was diminished, whereas pSer2 was refractory to the Cdk9 inhibitor at 20- to 100-fold higher doses (Fig. 1c). This is consistent with results in fission yeast, where Cdk9 is not a major contributor to pSer2 in vivo ^21,35^. Pol II pSer2 was also relatively refractory to Cdk9 depletion by short hairpin RNA (shRNA) in HCT116 cells ^9^. In fission yeast, chemical-genetic inhibition of Cdk9 led to rapid, nearly complete dephosphorylation of the Spt5 CTD (T_1/2_ ~20 sec); the rate of decay decreased ~4-fold in *dis2* mutant strains, suggesting that the fast kinetics in *dis2^+^* cells were partly due to the concomitant activation of Dis2 (PP1) when Cdk9 is inactivated ^27^. In HCT116 cells, both pThr806 and a phosphorylation outside the CTRs, pSer666, were lost rapidly upon treatment with 250 nM NVP-2 (T_1/2_ ~10 min), consistent with a similar, reinforcing effect of kinase inhibition and phosphatase activation (Fig. 1d).

To complete the potential Cdk9-PP1-Spt5 circuit in human cells, we sought to validate Spt5 as a target of PP1. Purified, recombinant PP1 was able to dephosphorylate a CTR1-derived peptide phosphorylated on the position equivalent to Thr806 in the intact protein, but was inert towards a pSer666-containing peptide derived from the KOW4-KOW5 linker (Fig. 1e). The pSer666 substrate was efficiently dephosphorylated by λ phosphatase, suggesting that this resistance was indeed due to restricted substrate specificity of PP1. We obtained similar results in assays of immunoprecipitated GFP-PP1 isoforms expressed in human cells (Supplementary Fig. 1b, c). Next we asked if Spt5 phosphorylation was sensitive to loss of PP1 function in vivo. Depletion of all three PP1 catalytic subunits with small interfering RNA (siRNA) increased steady-state levels of pThr806 in extracts (Fig. 1f), whereas knockdown of PP1 isoforms individually or in pairwise combinations had negligible effects on Spt5 phosphorylation (Supplementary Fig. 1d), suggesting redundancy or compensation. In contrast, even the triple knockdown had no effect on pSer666 (Fig. 1f, Supplementary Fig. 1e), consistent with the insensitivity of this modification to PP1 in vitro. Thus, 1) Cdk9 phosphorylates both Spt5 and PP1γ in vivo, 2) PP1 can dephosphorylate an Spt5 CTR1-derived peptide (but not a pSer666-containing peptide) in vitro, and 3) levels of pThr806 (but not pSer666) are limited by PP1 activity in vivo. Taken together, these results indicate that the enzymatic elements of an elongation-termination switch defined in fission yeast are conserved in human cells, but suggest that a different phosphatase might target pSer666 (and possibly other sites in the elongation complex), perhaps to support a different function.

### The Spt5 CTR is hypophosphorylated at the 3’ pause

We next asked if the *output* of Cdk9-PP1 signaling is similar in yeast and human cells. We first confirmed that antibodies against pThr806 ^9^ recognized CTR1 phosphorylated by Cdk9 in vitro (Supplementary Fig. 2a) but not unphosphorylated CTR1, or CTR2, another carboxy-terminal block of Thr-Pro-containing repeats in Spt5 ^36,37^. Reactivity with Spt5 overexpressed in human cells was diminished but not abolished by mutation of Thr806 to Ala (Supplementary Fig. 2b), suggesting that the antibody can recognize other repeats in CTR1. In ChIP-seq analysis in HCT116 cells (Supplementary Fig. 2c), the distribution of total Spt5 closely matched that of transcribing Pol II, with peaks near the TSS and downstream of the CPS, indicating promoter-proximal and 3’ pausing, respectively (Fig. 2a, b). This is consistent with the tight association of DSIF with Pol II in elongation complexes ^38,39^. In contrast, pThr806 (pCTR1) and pSer2—which have both been interpreted as markers of elongating Pol II—had different distributions. This was most evident downstream of the CPS, where total Spt5 accumulated together with Pol II; pSer2 peaked in this region, whereas pThr806 signals were diminished—a divergence evident both in metagene plots (Fig. 2a) and browser tracks from individual genes (Fig. 2b). Comparison of metagene plots of pThr806:Spt5 and pSer2:Pol II ratios revealed an inverse relationship (Fig. 2c): pThr806:Spt5 began to drop just upstream of the CPS and reached a minimum at the 3’ pause, whereas pSer2:Pol II increased at the CPS and peaked at the pause. The divergence between the two modifications at both the TSS and termination zone (TZ), despite the high correlation between total Pol II and Spt5, was confirmed by principal component analysis (Fig. 2d, Supplementary Fig. 2d, e).

**Fig. 2.**
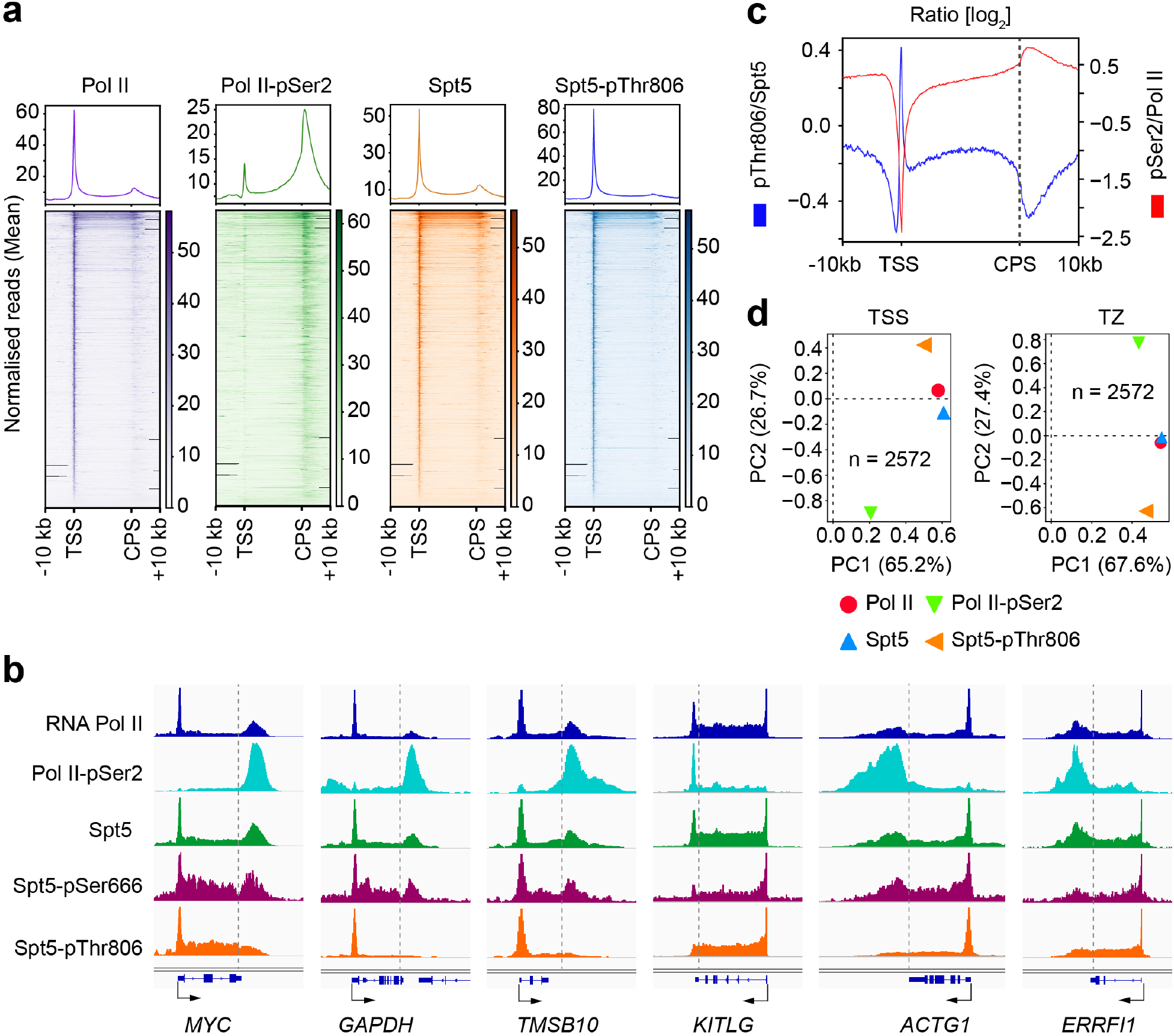
Spt5 CTR1 becomes dephosphorylated downstream of the CPS. **a** Metagene analyses and heat maps of ChIP-seq data (*n* = 20,130 genes) for total Pol II, pSer2, total Spt5 and Spt5-pThr806, as indicated (*n* = 2 biological replicates). **b** Individual gene tracks of Pol II, Pol II-pSer2, Spt5, Spt5-pSer666 and Spt5-pThr806. **c** Metagene analysis of pThr806:Spt5 and pSer2:Pol II ratios (*n* = 20,130 genes). **d** Principal component analysis of Spt5-Thr806 and Pol II-pSer2 around TSS (−50 to +500 bp) and termination zone (TZ; −50 to +5000 bp of the CPS) for *n* = 2,572 highly Pol II occupied genes (percentages in parentheses alongside axes indicate amount of variance explained).

A reduction in pCTR1 that precedes Pol II pausing and a pSer2 peak is consistent with a “sitting duck” model, whereby slowing of elongation triggers pSer2, recruitment of cleavage and polyadenylation factors and termination ^24,25^. We detected a similar, reciprocal relationship between Spt5-CTD phosphorylation and pSer2 in fission yeast ^27^, in which Pol II undergoes a metazoan-like pause downstream of the CPS ^40^. In both human and fission yeast cells, Cdk9 inhibition slows elongation in gene bodies ^21,22^; to ask if this had the predicted, opposite effects on pThr806 and pSer2, we treated HCT116 cells with NVP-2 and performed ChIP-qPCR analysis. A 1-hr treatment with 250 nM NVP-2 caused Pol II depletion from the gene body—as expected if promoter-proximal pause release was impeded—and near-complete loss of pThr806 on both *MYC* and *GAPDH* (Fig. 3a-c, Supplementary Fig. 3a-c). Absolute pSer2 levels were also diminished by NVP-2 treatment, but the pSer2:Pol II ratio was *increased* 2-3-fold in gene bodies, suggesting ectopically increased Ser2 phosphorylation due to slowed elongation.

**Fig. 3.**
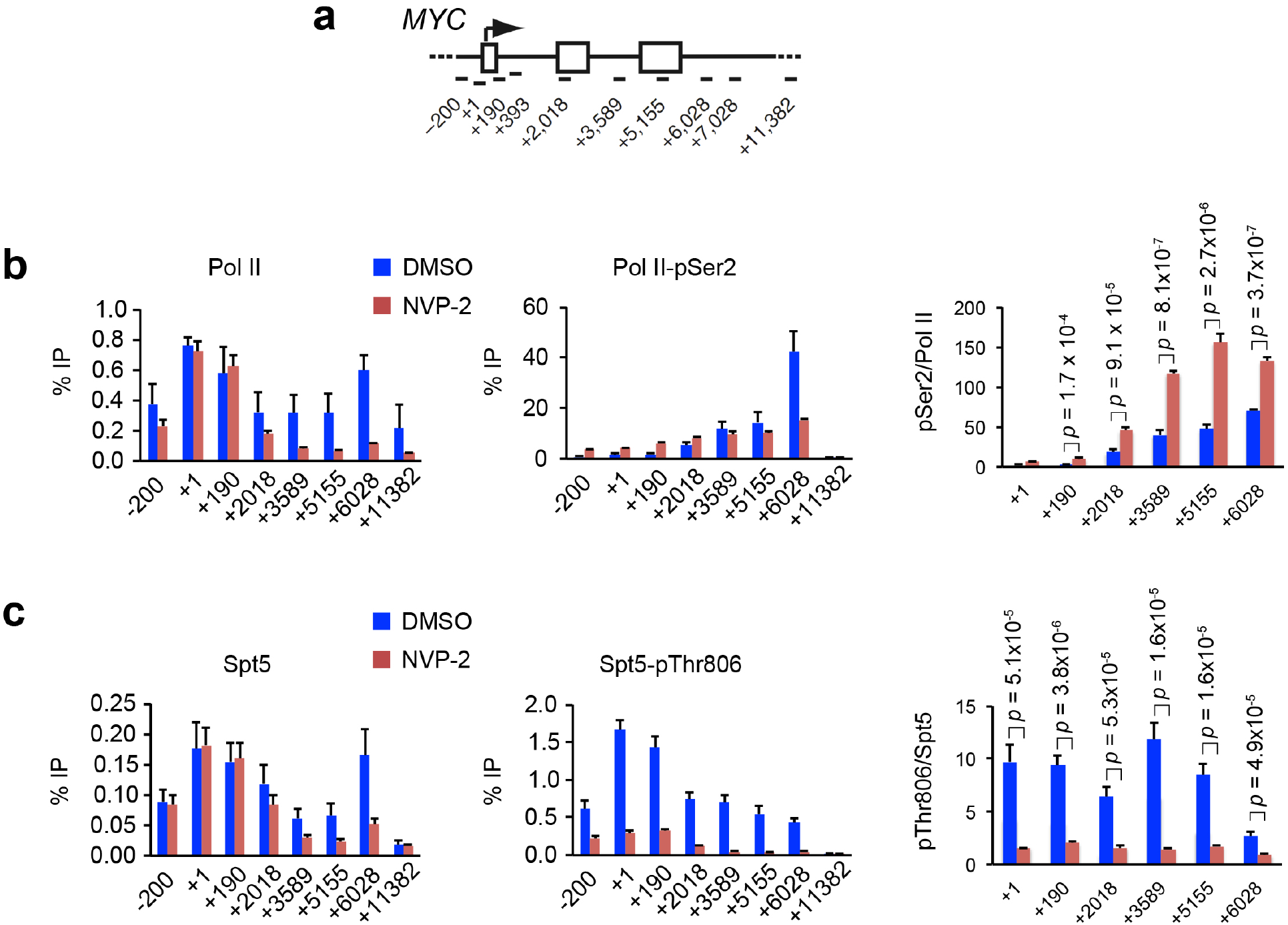
Cdk9 inhibition diminishes Spt5-CTR1 phosphorylation but increases Pol II CTD-Ser2 phosphorylation on chromatin. **a** Schematic of the *MYC* gene, indicating positions of primer pairs used in ChIP-qPCR analysis. **b** ChIP-qPCR analysis of total Pol II and CTD-Ser2 phosphorylation in HCT116 cells treated with NVP-2 (250 nM) or mock treated (DMSO) for 1 hr. **c** ChIP-qPCR analysis of total Spt5 and Spt5-Thr806 phosphorylation in HCT116 cells treated with NVP-2 (250 nM) or mock treated (DMSO) for 1 hr. Indicated *p* values were calculated using Student’s *t*-test. Error bars indicate + s.d. of four biological replicates (**b** and **c**).

### Spt5 phosphorylations are differentially distributed on chromatin

The metagene plots of pSer2:Pol II and pThr806:Spt5 ratios (Fig. 2c) also diverged at 5’ ends of genes, where the former had a deep trough—presumably reflecting pausing of Pol II with high Ser5 phosphorylation (pSer5) but low pSer2 ^2^—whereas the latter peaked, suggesting that paused complexes can contain high levels of pCTR1. Both pThr806 and pSer666 were among the many residues phosphorylated—in Spt5 and other components of the transcription machinery—when paused Pol II complexes were converted to elongating complexes by treatment with P-TEFb ^18^. We therefore asked if pSer666 was enriched in complexes that had escaped the pause. In contrast to the anti-pThr806 antibody, anti-pSer666 was unable to recognize CTR1 or CTR2, but did react with full-length Spt5, dependent on pre-incubation with Cdk9 and ATP (Supplementary Fig. 4a). Moreover, immunoblot reactivity of Spt5 in cell extracts required an intact Ser666 residue (Supplementary Fig. 4b), suggesting that the antibody is specific for a single modification, and possibly explaining the lower signals it produced, relative to anti-pThr806, in ChIP-seq analysis (Fig. 2b). Despite lower signals, pSer666 had a distribution similar to that of total Spt5, including a peak downstream of the CPS, where pThr806 decreased (Fig. 2b, 4a and b). A metagene analysis comparing pSer666:Spt5 and pThr806:Spt5 ratios revealed two differences: 1) a trough, rather than a peak, of pSer666:Spt5 near the TSS, followed by an increase upon entering the gene body; and 2) a shallower trough of pSer666:Spt5, relative to that of pThr806:Spt5, downstream of the CPS (Fig. 4c), suggesting that pSer666 was largely retained in the 3’-paused complex.

**Fig. 4.**
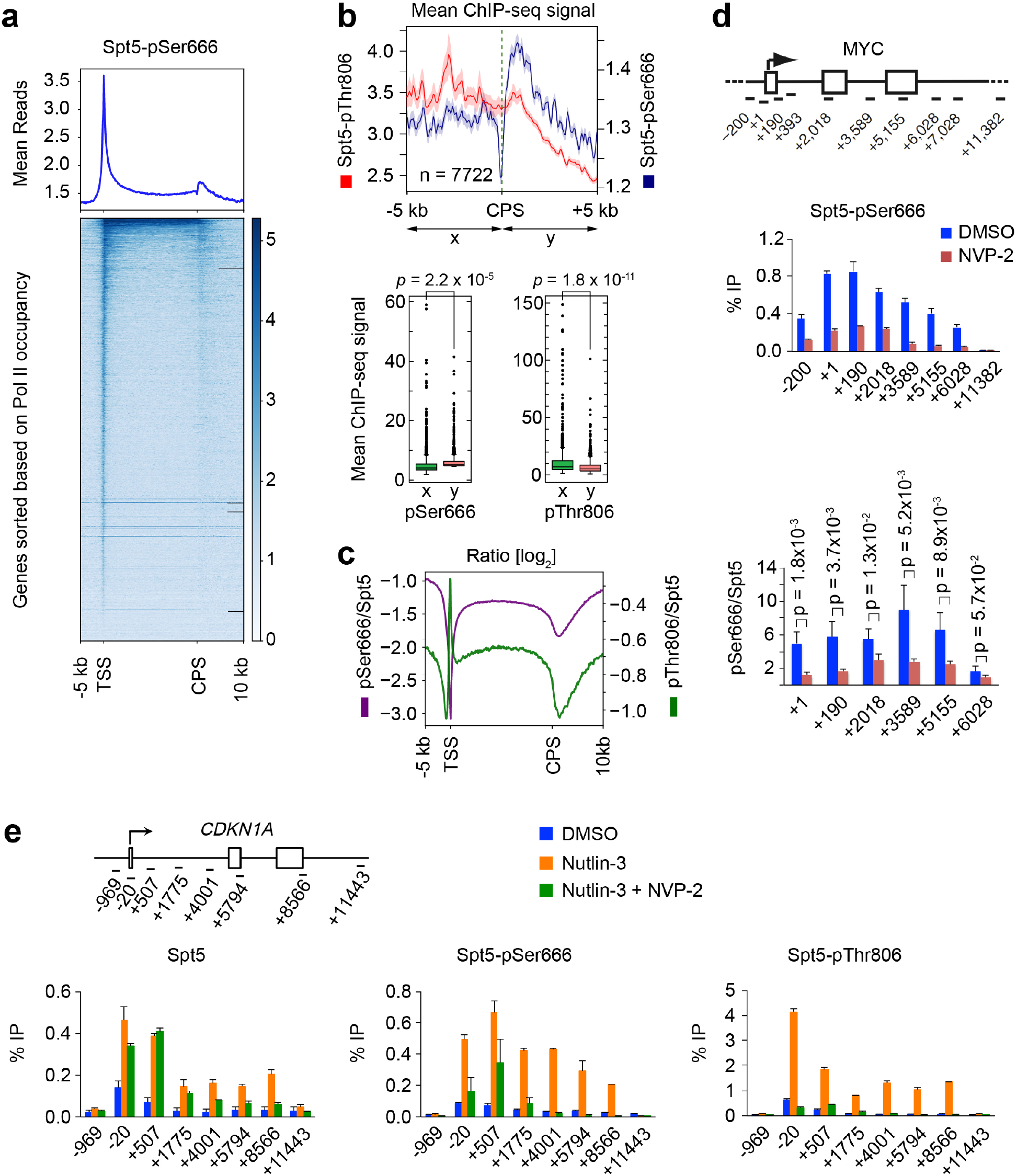
Chromatin distribution of Spt5-pSer666 is distinct from that of Spt5-pThr806. **a** Metagene analyses and heat maps (*n* = 20,130 genes) of ChIP-seq data for Spt5-pSer666 (*n* = 2 biological replicates). **b** Genes separated from their neighbors at both ends by >10 kilobases (*n* = 7,772) show significant accumulation of pSer666 but not pThr806 downstream of the CPS (top, metagene plots; bottom, box plots). **c** Metagene analysis of pSer666:Spt5 and pThr806:Spt5 ratios (*n* = 20,130 genes). **d** ChIP-qPCR analysis of Spt5-Ser666 phosphorylation in HCT116 cells treated with NVP-2 (250 nM) or mock treated (DMSO) for 1 hr. Indicated *p* values were calculated using Student’s *t*-test. Error bars indicate + s.d. of four biological replicates. **e** Schematic of the *CDKN1A* gene, indicating positions of primer pairs used in ChIP-qPCR analysis (top). ChIP-qPCR analysis of total Spt5, Spt5-pSer666 and Spt5-pThr806 in HCT116 cells mock treated (DMSO) or treated with 5 μM nutlin-3 alone or together with 250 nM NVP-2 for 2 hr. Error bars indicate + s.d. of four biological replicates.

The relative increases in pSer666 downstream of the TSS were also detectable by ChIP-qPCR analysis on *MYC* and *GAPDH* genes, and sensitive to Cdk9 inhibition (Fig. 4d, Supplementary Fig. 4c). To ask if transcriptional induction triggers increased pSer666, we performed ChIP-qPCR at the p53-responsive, pause-regulated *CDKN1A* gene ^41^. Upon p53 stabilization by nutlin-3 ^42^, Pol II was redistributed from the promoter-proximal pause site into the body of the *CDKN1A* gene (Supplementary Fig. 4d). Both total Spt5 and pThr806 signals increased roughly proportionally over the TSS and gene body, whereas pSer666 peaked ~0.5 kb downstream of the TSS (Fig. 4e). As was the case at constitutively expressed *MYC* and *GAPDH*, inhibiting Cdk9 diminished both pSer666 and pThr806 on the induced *CDKN1A* gene, but increased the pSer2:Pol II ratio in the ~2-kb region downstream of the TSS (Fig. 4e and Supplementary Fig. 4d).

ChIP-seq analysis comparing nutlin-3- to mock-treated HCT116 cells revealed differential distributions of pSer666 and pThr806 on p53-responsive genes. Browser tracks of two representative p53 targets, *CDKN1A* and *GDF15* (Fig. 5a), as well as metagene plots of nutlin-3-induced genes (Fig. 5b, Supplementary Fig. 5a), showed 1) increased pSer666 but not pThr806, relative to total Spt5, in the region just downstream of the TSS; and 2) retention of pSer666 but diminution of pThr806 at the 3’ pause downstream of the CPS, where pSer2 signals were maximal. Quantification of reads downstream of the TSS revealed more significant increases in pSer666 than in pThr806 or pSer2, with the largest gains occurring on non-pause-regulated p53 target genes (Fig. 5c, Supplementary Fig. 5b-d). Downstream of the CPS there was a significant increase in pSer666 but not pThr806 in response to p53 induction (Supplementary Fig. 5e). Given the dependence of both pThr806 and pSer666 on Cdk9, their differential distributions might reflect removal by different phosphatases, a possibility we explore in the next section.

**Fig. 5.**
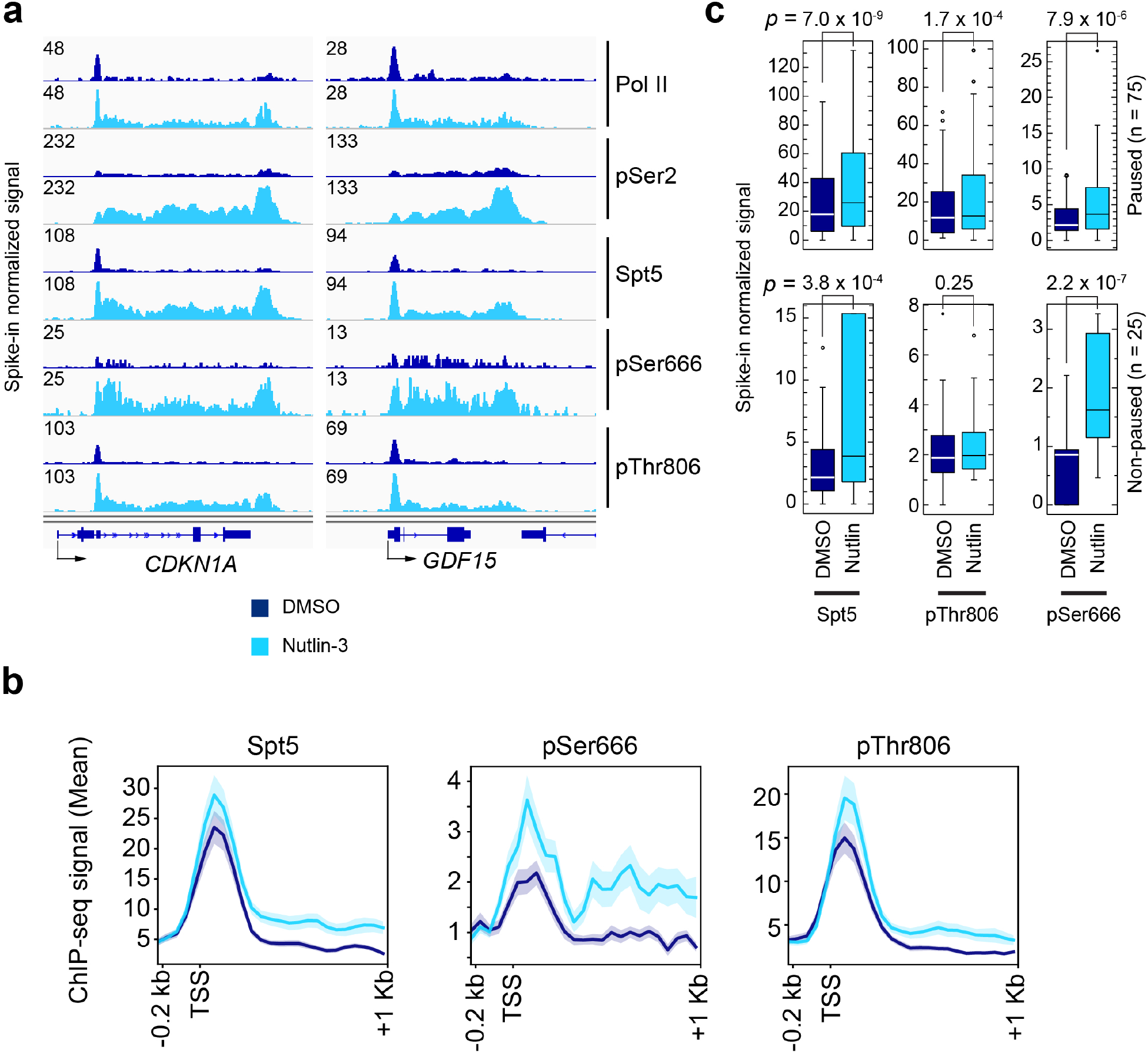
Phosphorylation of Spt5-Ser666 is increased downstream of the TSS and retained downstream of the CPS on genes induced by p53 activation. **a** Individual ChIP-seq gene tracks at *CDKN1A* and *GDF15*—two p53 targets—in HCT116 cells mock-treated (DMSO) or treated with 5 μM nutlin-3 for 2 hr. **b** Metagene analysis of Spt5, Spt5-pThr806 and Spt5-pSer666 at genes induced by nutlin-3 (*n* = 75; paused genes). **c** Box plots comparing ChIP-seq reads in first 100-nt interval downstream of TSS, in the absence or presence of nutlin-3, for Spt5, Spt5-pThr806 and Spt5-pSer666 at genes induced by nutlin-3, divided into those classified as pause-regulated (pause index ≥ 2.0, *n* = 75) or non-paused (pause index < 2.0, *n* = 25).

### PP4 is a Cdk9-regulated Spt5 phosphatase that supports promoter-proximal pausing

We reasoned that a pSer666 phosphatase might also be a target of negative regulation by Cdk9, based on the similar kinetics of pThr806 and pSer666 dephosphorylation after Cdk9 inhibition (Fig. 1d). Among the sites labeled by Cdk9 in HCT116 whole-cell extracts was Thr173 of the PP4 regulatory subunit PP4R2 ^9^. This residue and several others in the PP4 complex were previously shown to be phosphorylated, by a CDK, to inhibit PP4 activity in response to mitotic-spindle poisons ^43^. An anti-phospho-Thr173 antibody failed to detect this modification in cell extracts or in untreated anti-PP4R2 immunoprecipitates, but recognized immobilized PP4R2 after treatment with Cdk9 in vitro, validating PP4R2 as a potential P-TEFb substrate (Fig. 6a). A phosphatase precipitated with either anti-PP4R2 or anti-PP4C (catalytic subunit) antibodies was active towards both pSer666- and pThr806-containing phosphopeptides (Fig. 6b), in contrast to PP1, which only worked on the latter (Fig. 1e). Pre-treatment of HCT116 cells with either NVP-2 or RO-3306 increased the phosphatase activity of anti-PP4R2 or −PP4C immunoprecipitates (Fig. 6b) without affecting immunoprecipitation efficiency or complex integrity (Supplementary Fig. 6a), suggesting negative regulation of PP4 activity by Cdk9 or Cdk1—similar to the situation with PP1γ (Fig. 1b). Moreover, incubation of anti-PP4R2 immunoprecipitates with purified Cdk9 and ATP prior to a phosphatase assay reduced activity ~3-fold, indicating that PP4 complexes were sensitive to direct inhibition by P-TEFb (Fig. 6c).

**Fig. 6.**
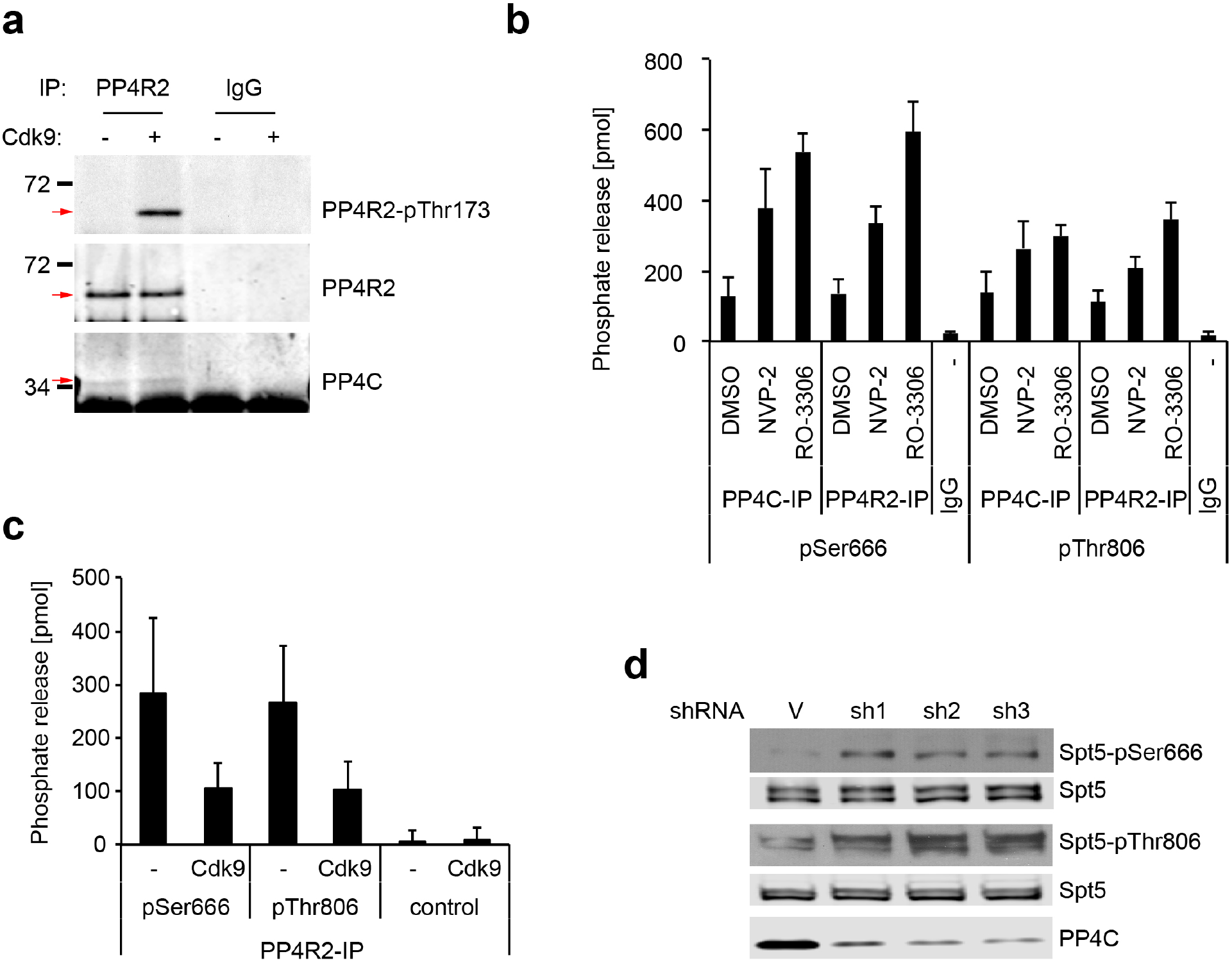
PP4 is a potential Spt5 phosphatase, active towards pSer666 and pThr806 and subject to negative regulation by Cdk9. **a** PP4R2 was immunoprecipitated from HCT116 cell extracts, incubated in vitro with purified, recombinant Cdk9/cyclin T1 and ATP, and immunoblotted with antibodies to PP4R2-pThr173, total PP4R2 or PP4C, as indicated at *right*. **b** HCT116 cells were mock-treated (DMSO) or treated with inhibitors of Cdk9 (NVP-2) or Cdk1 (RO-3306), as indicated, for 1 hr before lysis and extract preparation. Anti-PP4R2, anti-PP4C or control IgG immunoprecipitates, as indicated, were incubated with Spt5-derived phosphopeptides containing pSer666 or pThr806, as indicated, and phosphate release was measured colorimetrically. Error bars indicate + s.d. of six biological replicates. **c** Anti-PP4R2 immunoprecipitates were incubated with 5 ng purified, recombinant Cdk9/cyclin T1 and ATP or mock-treated (as indicated), washed and tested for phosphatase activity towards an Spt5-derived phosphopeptide containing pSer666, as in **b**. Error bars indicate + s.d. from three biological replicates. **d** HCT116 cells were infected with lentivirus expressing shRNA targeting PP4C (three different hairpins) or a non-targeted control vector (V), and chromatin fractions were immunoblotted for Spt5-pSer666, Spt5-pThr806, total Spt5 (to ensure equal loading) and total PP4C (to assess efficiency of depletion).

To test whether PP4 regulated Spt5 phosphorylation in vivo, we depleted PP4 by infection with lentivirus vectors expressing shRNA targeting PP4C. Three different shRNAs each diminished PP4C levels by ~70-80%, and increased pSer666:Spt5 and pThr806:Spt5 signal ratios in immunoblots of chromatin extracts (Fig. 6d, Supplementary Fig. 6b). This was in contrast to effects of PP1 depletion, which preferentially affected pThr806 levels (Fig. 1f), but consistent with the substrate specificity of immunoprecipitated PP4 complexes in vitro (Fig. 6b).

To ask if differential localization of PP4 and PP1 might contribute to the different spatial distributions of Spt5 phospho-isoforms on chromatin, we performed ChIP-qPCR analysis of PP4C, PP4R2 and PP1γ on the *MYC*, *GAPDH* and *CDKN1A* genes (Fig. 7a, b and Supplementary Fig. 7a). Both PP4 subunits crosslinked predominantly between the TSS and ~2-3 kb downstream, and were present at low or undetectable levels near the 3’ ends of genes. PP1γ had nearly the opposite distribution, crosslinking at near-background levels between the TSS and +2 kb before peaking close to the CPS, consistent with its residence in the CPF ^28,44,45^. We also performed ChIP-qPCR analysis in cells exposed to NVP-2 (250 nM, 1 hr); this treatment had minimal effects on chromatin association of total PP4C (Supplementary Fig. 7b) or PP4R2 (Fig. 7c, left panel), but decreased the signals obtained with the anti-phospho-PP4R2-Thr173 antibody to near-baseline levels (Fig. 7c, middle and right panels), consistent with negative regulation of PP4 by P-TEFb on chromatin.

**Fig. 7.**
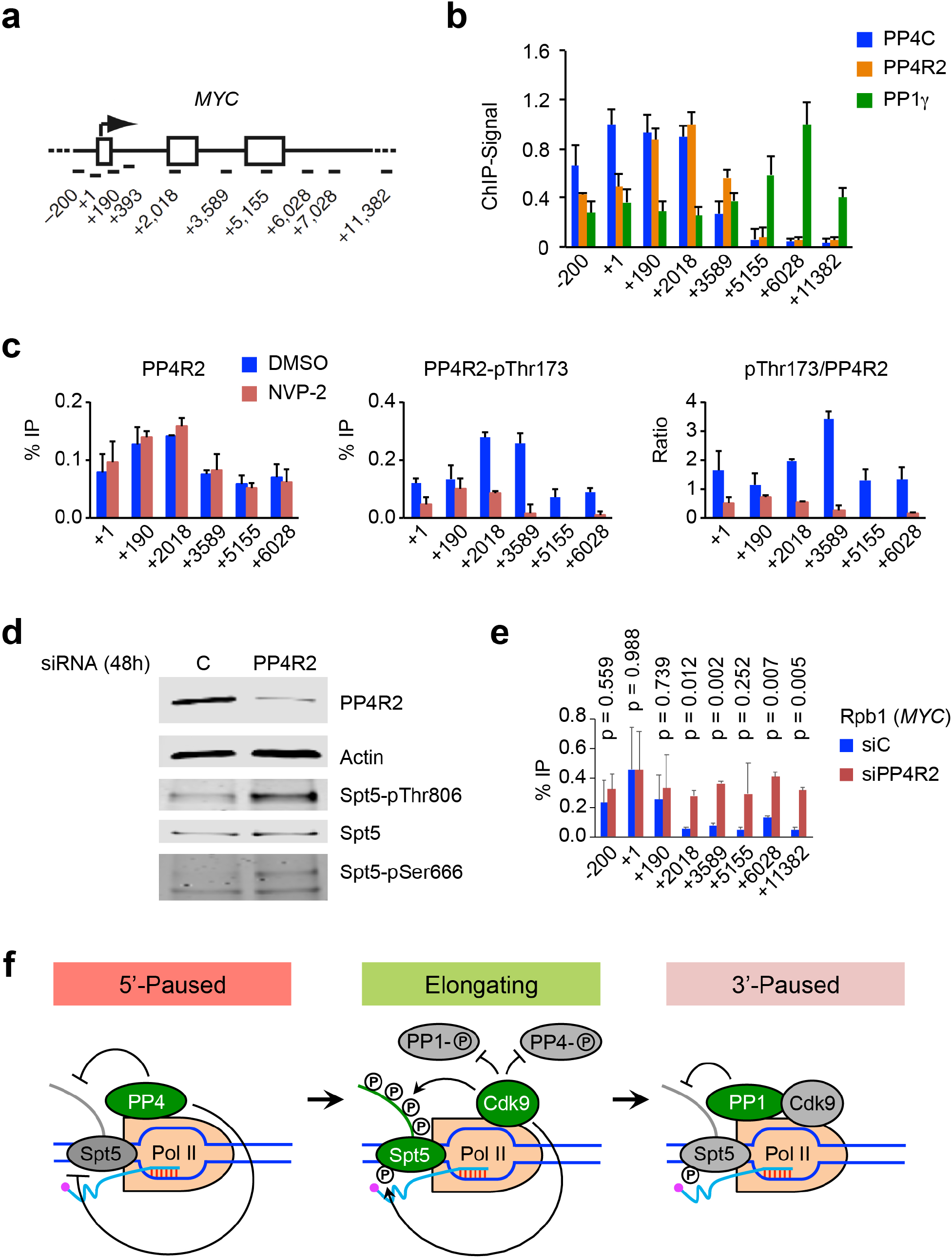
Distinct spatial distributions and specificities of two Cdk9-regulated phosphatases serve to order the transcription cycle. **a** Schematic of the *MYC* gene, indicating positions of primer pairs used in ChIP-qPCR analysis. **b** ChIP-qPCR analysis of PP4C, PP4R2 and PP1γ on the *MYC* gene in unperturbed HCT116 cells. Error bars indicate + s.d. of two biological replicates. **c** ChIP-qPCR analysis of PP4R2 and PP4R2-pThr173 after inhibition of Cdk9 with NVP-2 (250 nM) or mock treatment (DMSO) for 1 hr. Error bars indicate + s.d. of two biological replicates. **d** Cells transfected with siRNA targeting PP4R2 or non-targeted negative control siRNA were subjected to formaldehyde crosslinking, chromatin isolation and reversal of crosslinking before analysis by immunoblotting with indicated antibodies to measure the depletion of PP4R2 and phosphorylation of Spt5 at Ser666 and Thr806. **e** ChIP-qPCR analysis on *MYC* shows Pol II distribution with or without PP4R2 depletion. Error bars (+ s.d.) and *p*-values were calculated using data from four biological replicates (*n* = 4). **f** Two distinct Cdk9-phosphatase switches govern transitions in Spt5 phosphorylation state—a model. At the 5’ pause prior to P-TEFb activation, the PP4 complex is active and both CTR1 and the KOW4-KOW5 loop are unphosphorylated, At the 3’ pause, PP1 becomes active to dephosphorylate CTR1 but not Ser666, distinguishing the two paused complexes. During elongation, Cdk9 phosphorylates Spt5, and inhibits PP4 and PP1.

Finally, we asked if we could mimic effects of P-TEFb activity on Pol II distribution by decreasing cellular levels of PP4R2. Depletion of PP4R2 with small interfering RNA (siRNA) nearly abolished both total PP4R2 and PP4R2-pThr173 ChIP signals (Supplementary Fig. 7c), increased both pSer666 and pT806 in extracts (Fig. 7d) and shifted the distribution of Pol II into the bodies of the pause-regulated *MYC*, *GAPDH* and *CDKN1A* genes (Fig. 7e, Supplementary Fig. 7d-f). This suggests a role of PP4 in imposing a barrier to elongation at the promoter-proximal pause; this barrier can apparently be lowered artificially by depletion of PP4R2 or, we surmise, physiologically by PP4-inhibitory phosphorylation catalyzed by Cdk9. Taken together, our results suggest that two distinct Cdk9-phosphatase circuits operate at the beginning and end of the elongation phase in the Pol II transcription cycle (Fig. 7f).

## Discussion

Spt5 is an ancient component of the transcription machinery, with functions in elongation and termination conserved in eukaryotes, archaea and prokaryotes ^46^. Although studies of metazoan DSIF have emphasized its central roles in promoter-proximal pausing, recent work in yeast indicates functions for the Spt4/Spt5 heterodimer throughout the elongation phase and during termination ^47,48^. Structural analyses reveal tight association between DSIF and the clamp region of Pol II in elongation complexes assembled from purified components ^38,39^. Pol II and Spt5 can be cross-linked co-extensively along the entire lengths of genes ^27,47,48^, also consistent with Spt5 acting throughout the transcription cycle.

A paused complex reconstituted in vitro with purified Pol II, DSIF and NELF was converted to an activated elongation complex by addition of elongation factors PAF and Spt6 and phosphorylation by P-TEFb ^18,19^. The activated complex contained numerous sites phosphorylated by Cdk9 on Rpb1 (in both the CTD and the linker connecting the CTD to the rest of the protein), Spt5, Spt6 and multiple subunits of the PAF complex.

There were 14 phosphorylations on Spt5 alone, including 1) pThr806 and several other CDK-consensus (Thr-Pro) motifs in CTR1, and 2) pSer666 and two other Ser residues in the KOW4-KOW5 linker. These phosphorylations are likely to be reinforcing, such that modification of any individual site—or even whole domains—may not be necessary to promote release from the promoter-proximal pause. To circumvent potential redundancy of modifications within Spt5 and other components of the elongation complex, we focused on defining roles for the relevant modifying enzymes. Our results indicate that two classes of Spt5 mark placed by Cdk9 near the beginning of the transcription cycle are removed at different times, by different phosphatases. This raises the possibility that they perform different roles in regulating either intrinsic properties of the elongation complex (e.g. catalytic rate) or its interactions with other factors—evidence of an Spt5-phosphorylation “code.” Although both pSer666 and pThr806 are detectable over much of the gene body, pThr806 is enriched at the promoter-proximal pause whereas the pSer666:Spt5 ratio increases further downstream, suggesting that KOW4-KOW5 linker phosphorylation is more likely to occur at or after pause release. Downstream of the CPS, where elongation is slowed, pCTR1 drops—similar to the pattern observed in fission yeast ^27^—whereas pSer666 is retained. This difference can be explained, without invoking additional kinases, by the inability of CPF-associated PP1 to dephosphorylate pSer666.

A decisive role in termination for PP1 and the Spt5 CTRs is supported by genetic interactions in fission yeast. Mutations of *spt5* that prevent CTD repeat phosphorylation at position Thr1 (*spt5-T1A*) suppressed a conditional *dis2-11* mutation, and mimicked termination-promoting effects of allele-specific Cdk9 inhibition ^21,27,29^. The narrowing of the termination zone by genetic manipulations in yeast was similar to the effects of introducing an intrinsically slow Pol II mutant variant or a Cdk9 inhibitor in human cells ^49,50^. Conversely, *dis2* loss-of-function alleles broadened the termination zone ^27,29^, as did a fast variant of human Pol II ^49^. These results supported the idea that Spt5-CTR phosphorylation by Cdk9 is an accelerator of elongation ^21,22^, whereas reversal of that phosphorylation by PP1 is a brake. Two recent reports extended this model to human cells, by showing that PP1 promotes Pol II slowing and termination through Spt5 dephosphorylation ^30,31^. Another implicated PP4 in regulation of Spt5 phosphorylation and function during early stages of transcription in *Caenorhabditis elegans* ^51^. The results presented here establish Cdk9 as the linchpin of this network, able to phosphorylate multiple domains of Spt5 while restraining the activity of both PP4 and PP1 through inhibitory phosphorylation.

Based on our results we propose a phosphorylation-dephosphorylation cycle during transcription elongation, governed by Cdk9 and (at least) two opposing phosphatases, PP4 and PP1 (Fig. 7f). Localization of PP4 to 5’ gene regions would help stabilize the promoter-proximal pause by keeping CTR1 and Ser666 (and possibly other sites in the paused complex) unphosphorylated until Cdk9 is recruited and activated. During pause release and subsequent elongation, Cdk9 reinforces its phosphorylation of Spt5 in both CTR1 and the KOW4-KOW5 loop by inhibiting both PP4 and PP1. As transcription complexes traverse the CPS there is a switch from a high-Cdk9/ low-PP1 state to its opposite—and thus from high to low CTR1 phosphorylation—by an undetermined mechanism. Exclusion of PP4 from downstream regions would ensure that PP1-resistant marks such as pSer666 persist in the 3’-paused complex, possibly to distinguish it from the complex paused in the promoter-proximal region.

The model accounts for differential distributions of pSer666, pThr806 and pSer2 on chromatin, and the biochemical relationships underpinning it have been validated by results of kinase and phosphatase assays in vitro and of enzyme inhibition or depletion in vivo. Selective inhibition of Cdk9 in human cells stimulated the phosphatase activity of PP4 complexes and diminished PP4R2-pThr173 signals on chromatin. Cdk9 inhibition also diminished phosphorylation of a known inhibitory site on PP1γ in extracts, although we were unable to detect this modification on chromatin with available phosphospecific antibodies. Finally, our studies do not rule out contributions by other kinases and phosphatases, possibly arranged in similar switch-like circuits, to the regulation of Spt5 phosphorylation or Pol II elongation.

We describe two regulatory circuits involving Cdk9 and distinct phosphatases subject to inhibitory phosphorylation by Cdk9. Both PP1 and PP4 are also inactivated by cell-cycle CDKs that phosphorylate either the PP1 catalytic subunit ^34,52,53^ or PP4R2 ^43^. Cdk1 inhibits PP1 during mitosis; a drop in Cdk1 activity due to cyclin B degradation at metaphase leads to PP1 activation, dephosphorylation of mitotic phosphoproteins and mitotic exit ^34,54^. We proposed that an analogous mechanism controls “transcription exit” through PP1-dependent Spt5 dephosphorylation in fission yeast ^27^; here we provide evidence for conservation of this mechanism. A similar interaction between PP4 and a cell-cycle CDK may regulate the nucleation of microtubules at centrosomes ^43^; we now implicate PP4 in a Cdk9-containing circuit regulating the transition to processive elongation.

Therefore, analogous modules, consisting of a CDK and an opposing phosphatase that is also a CDK substrate, govern transitions in both transcription and cell-division cycles. One property conferred by this arrangement is switch-like, all-or-none behavior: CDK inactivation causes rapid target dephosphorylation because it simultaneously activates the relevant phosphatase. We envision that transitions in the transcription cycle, such as changes in elongation rate dictated by template sequence elements or chromatin features, are naturally switch-like. Moreover, linking activity of a single CDK to phosphatases with different specificities can generate spatially diverse patterns, even for modifications within a single effector protein. We note that the Cdk9-phosphatase circuits described here impart positional information, differentiating a promoter-proximally paused complex (pSer666 OFF/ pCTR1 OFF) from one paused at the 3’ end (pSer666 ON/ pCTR1 OFF). This is a fundamental principle of cell-division control; one cyclin-CDK complex can drive the entire cell cycle, producing myriad different temporal patterns of target protein phosphorylation and function, in part through the action of multiple phosphatases ^55^. We propose this as a strategy to order the Pol II cycle, in which relatively few CDKs (<10) are needed to phosphorylate hundreds of substrates that act at different steps of transcription ^7,9^.

## Methods

### Cell lines and drug treatments

Colon carcinoma-derived HCT116 cells were cultured in McCoy’s 5A medium with L-glutamine (Corning) supplemented with 10% Fetal Bovine Serum (FBS, Gibco) and 1x Penicillin-Streptomycin (Corning). Drug treatments were performed at 50-60% confluence by substituting the growth medium with fresh medium containing either DMSO, 250 nM (except where noted) NVP-2 (provided by N.S. Gray), 10 μM RO-3306 (Selleckchem) and 5 μM nutlin-3 (Cayman Chemical Company).

### Antibodies

The antibodies used were: rabbit anti-Rpb1 (sc-899; Santa Cruz Biotechnology), rabbit anti-Rpb1 (A304-405A & A304-405A, Bethyl Laboratories), rabbit anti-Rpb1 CTD pSer2 (ab5095, Abcam), rabbit anti-Spt5 (A300-868A, Bethyl Laboratories), mouse anti-Spt5 (sc-133217, Santa Cruz Biotechnology), rabbit anti-Spt5-pSer666 and −pThr806 (21^st^ Century Biochemicals) previously described ^56^, rabbit anti-PP4R2 (A300-838A, Bethyl Laboratories), rabbit anti-PPP4C (A300-835A, Bethyl Laboratories), sheep anti-PP4R2-pThr173 (Division of Signal Transduction Therapy, University of Dundee Scotland) ^43^, rabbit phospho-PP1α (Thr320) antibody (2581S, Cell Signaling Technology), mouse anti-GFP (sc-9996, Santa Cruz Biotechnology), mouse anti-pan PP1 (sc-7482, Santa Cruz Biotechnology), goat anti-PP1α (sc-6104, Santa Cruz Biotechnology), mouse anti-PP1β (sc-373782, Santa Cruz Biotechnology), rabbit anti-PP1γ (A300-906A, Bethyl Laboratories), goat anti-PP1γ (sc-6108, Santa Cruz Biotechnology), mouse anti-tubulin (T5168, Sigma-Aldrich), rabbit anti-GST (sc-459, Santa Cruz Biotechnology), mouse anti-FLAG® M2 (F3165, Sigma-Aldrich) and rabbit anti-FLAG (2368, Cell Signaling).

### Protein Extraction

Whole-cell extracts were prepared as follows: Cells were washed twice with cold phosphate-buffered saline (PBS) and collected in RIPA buffer (50 mM Tris-HCl at pH 8.0, 150 mM NaCl, 1% nonidet-P-40, 0.5% sodium deoxycholate, 0.1% sodium dodecyl sulfate) supplemented with protease inhibitors (10 μM Pepstatin A, 1 μM Leupeptin, 2 mM 4-(2-aminoethyl)benzenesulfonyl fluoride hydrochloride [AEBSF], 1 μM Aprotinin, 1mM phenylmethylsulfonyl fluoride [PMSF]), phosphatase inhibitors (40 mM sodium β-glycerophosphate, 4 mM Na_3_VO_4_, 50 mM NaF) and 1 mM DTT. Cells were lysed in a Bioruptor (Diagenode) for 10 min with cycles of 30 sec ON and 30 sec OFF. The lysate was clarified by centrifugation at 4°C at 20,000 × g_av_ for 10 min. The chromatin fraction was prepared as described previously ^57^.

### RNAi

PP1 isoform-specific siRNAs targeting PP1α (sc-36299), PP1β (sc-36295) and PP1γ (sc-36297) were from Santa Cruz Biotechnology. For pan-PP1 depletion equimolar concentrations of each siRNA were used. HCT116 cells were transfected using Lipofectamine RNAiMAX (Invitrogen) according to manufacturer’s instructions. Cells were collected 24 hr post-transfection for lysate preparation and immunoblotting. Human embryonic kidney (HEK293) cells cultured in Dulbecco′s Modified Eagle′s Medium (DMEM), supplemented with 10% FBS and 1x Penicillin-Streptomycin, were used to generate lentivirus particles expressing PP4C-targeting shRNA obtained from Sigma: shRNA-1 (TRCN0000010737), shRNA-2 (TRCN0000272746), and shRNA-3 (TRCN0000272747). HCT116 cells were infected with pLKO.1-puro shRNA lentivirus and selected for 96 hr with 2 μg/ml puromycin and protein depletion verified by immunoblot. For PP4R2 depletion, siRNA targeting PP4R2 (sc-78526, Santa Cruz Biotechnology) was used. HCT116 cells were transfected using Lipofectamine RNAiMAX according to the manufacturer’s instructions. After 48 hr of transfection, cells were crosslinked with 1% formaldehyde for ChIP.

### Mutagenesis and ectopic protein expression

Phosphorylated residues of Spt5 were substituted with Ala or Asp using site-directed mutagenesis kits (Agilent Technologies), the oligonucleotides listed in Supplementary Table 1, and pCDNA-N-FLAG-SUPT5H (Sino Biological) as a template, according to manufacturer’s protocols. HCT116 cells were transfected with pCDNA-N-FLAG-SUPT5H-variants using Lipofectamine 3000 (Invitrogen) according to manufacturer’s instructions. Whole-cell lysates were immunoprecipitated with mouse anti-FLAG antibody, and immunoblotted with anti-pSer666, anti-pThr806, anti-Spt5 or rabbit anti-FLAG.

### Immunoprecipitation and immunoblotting

To immunoprecipitate proteins, 2 mg of whole-cell protein extract was incubated with antibodies for 4 hr at 4°C in RIPA buffer. Protein G Sepharose^TM^ 4 Fast Flow (GE Healthcare), pre-blocked with 1 mg/ml bovine serum albumin (BSA, Gemini Bio-products) and 0.25 mg/ml salmon sperm DNA (Trevigen) was added, and the resulting suspension was incubated for 2 hr at 25°C.

Beads were washed three times with ice-cold RIPA buffer. For immunoblot analysis, proteins were separated by SDS-PAGE and transferred to Amersham^TM^ Protran^TM^ 0.45 μm nitrocellulose membranes (GE Healthcare Life Sciences). The membranes were probed with primary antibodies at dilutions recommended by the suppliers. PP4R2 phospho-specific antibody (anti-PP4R2-pThr173) was used in the presence of a 10-fold molar excess of the appropriate non-phosphorylated peptide. Immunoblots were developed with either Horseradish Peroxidase (HRP) conjugated donkey anti-rabbit (NA934V, GE Healthcare Life Sciences), sheep anti-mouse (NA9310V, GE Healthcare Life Sciences), donkey anti-sheep (713-035-147, Jackson ImmunoResearch), donkey anti-goat (sc-2020, Santa Cruz Biotechnology), or Alexa Fluor-coupled goat anti-rabbit (A21076, Life Technologies), goat anti-mouse (A11375, Life Technologies), or donkey anti-goat (705-625-147, Jackson ImmunoResearch). Proteins were detected either by enhanced chemiluminescence (ECL, HyGLO HRP detection kit, Denville Scientific) or with Odyssey Imaging System (LI-COR Biosciences).

### Kinase and phosphatase assays

To detect Cdk9-dependent phosphorylations, GFP-PP1γ or PP4R2 were immunoprecipitated from whole-cell extracts. The bead-bound proteins were subjected to Cdk9 treatment as described ^15^. Briefly, immunoprecipitated proteins were either mock–treated or treated with purified Cdk9/cyclin T1 (5-10 ng) and ATP (1 mM) in kinase assay buffer (25 mM HEPES, pH 7.4, 10 mM NaCl, 10 mM MgCl_2,_ 1 mM DTT) for 30 min at 25°C. The beads were washed three times with RIPA buffer and analyzed by SDS-PAGE and immunoblotting. The PP4R2 samples (mock- and Cdk9-treated) were divided into two equal parts for immunoblot analysis and phosphatase assays. To measure protein phosphatase activity, GFP-tagged PP1 isoforms (PP1α, PP1β and PP1γ), PP4C and PP4R2 immunoprecipitated from whole-cell extracts were incubated with 50 μM phosphopeptide (Spt5-pThr806, Spt5-pSer666, H3pSer10) at 37°C for 1 hr. Colorimetric assays were performed in triplicate using BioMOL Green (Enzo Life Sciences) in 25 mM HEPES (pH 7.5), 100 mM NaCl, 1 mM MnCl_2,_ 1 mM DTT, in 96-well plates as described in manufacturer’s protocol. To test specificity of anti-pSer666 and anti-pThr806 antibodies, purified Cdk9/cyclin T1 was used to phosphorylate substrates purified from *E. coli* expressing DSIF (Spt4/Spt5 heterodimer), GST-CTR1 (amino acids 720-830 of Spt5 fused to glutathione-*S*-transferase) and GST-CTR2 (amino acids 844-1087 of Spt5) at a kinase:substrate ratio of 1:2000 for 15 min at 25°C in kinase assay buffer. The reactions were analyzed by SDS-PAGE and immunoblotting with anti-pSer666, anti-pThr806, anti-GST or anti-Spt5.

### ChIP-qPCR

ChIP-qPCR experiments were done as described previously ^15^. In brief, HCT116 cells grown to 50-60% confluence were crosslinked with 1% formaldehyde (Fisher Scientific) for 10 min at 25°C. Crosslinking was quenched with 125 mM glycine for 5 min at 25°C. Cells were washed twice with ice-cold PBS and collected into 1 ml of RIPA buffer supplemented with protease and phosphatase inhibitors for each 150-mm dish. Cells were lysed and chromatin sheared by sonication in a Bioruptor at high power, for 5 × 10 min with cycles of 30 sec ON and 30 sec OFF. Lysates were clarified by centrifugation at 20,000 × g_av_ for 20 min at 4°C. Before immunoprecipitation, lysates (~5 × 10^6^ cells per experiment) were pre-cleared with Pierce^TM^ Protein A Agarose (Thermo Scientific) for 2 hr at 4°C. Beads were separated by centrifugation at 4000 × g_av_ for 1 min at 4°C. The resulting supernatant was incubated with antibodies for 4 hr at 4°C with constant nutation. The suspension was incubated at 25°C for an additional 2 hr with Protein G Sepharose^TM^ 4 Fast Flow or Dynabeads^TM^ Protein G (Invitrogen), pre-blocked with 1 mg/ml BSA and 0.25 mg/ml Salmon Sperm DNA. The beads were washed with 2 × RIPA buffer, 4 × Szak IP wash buffer (100 mM Tris-HCl, pH 8.5, 500 mM lithium chloride, 1% (v/v) nonidet-P-40, 1% (w/v) sodium deoxycholate), 2 × RIPA buffer and 2 × TE buffer. After all wash steps, centrifugation was performed at 1700 × g for 1 min at 4°C. Protein-nucleic acid complexes were eluted from beads with elution buffer (46 mM TrisHCl, pH 8.0, 0.65 mM EDTA, 1% SDS) by incubating at 65°C for 15 min with occasional vortexing. Reversal of crosslinking was done by incubating at 65°C for 16 hr. The un-crosslinked suspension was treated with 1 μg of RNase A at 37°C for 30 min and with 0.8 units of Proteinase K (NEB) at 45°C for 45 min. DNA was purified using QIAquick^®^ PCR Purification Kit (Qiagen) according to the manufacturer’s protocol. The purified DNA was subjected to either qPCR with Radiant^TM^ Green qPCR Master Mix (2x) (Radiant Molecular Tools) in 386-well plates, or library preparation for sequencing.

### ChIP-seq

Preparation of multiplexed ChIP-seq libraries from purified immunoprecipitated chromatin and sequencing were performed as described ^27^. In brief, the NEBNext Ultra II DNA Library Preparation kit was used to generate libraries using 5-10 ng of input or immunoprecipitated DNA and barcode adaptors (NEBNext Multiplex Oligos for Illumina (Set 1, E7335 and Set 2, E7500)). Single-end (75-nt reads) or paired-end (40-nt reads) sequencing was performed on an Illumina NextSeq 500.

### Bioinformatic and statistical analysis

We used ‘FastQC Read Quality Reports’ (Galaxy Version 0.72) and ‘Trimmomatic Flexible Read Trimming Tool’ (Galaxy Version 0.36.6) to check quality of the sequencing reads and for barcode trimming, respectively. Trimmed sequencing reads were aligned to the human genome (version b37, hg19) using Bowtie2 ^58^ in Galaxy (Galaxy Version 2.3.4.2). Normalization of the aligned reads was done using ‘bamCoverage’ (Galaxy Version 3.1.2.0.0) by 1) computing and applying scaling factor obtained using aligned sequencing reads of the spike-in reference genome (for spike-in samples) and 2) by computing RPKM (reads per kilobase per million) (for the samples without spike-in control). Aligned sequences of each biological replicate were processed separately to identify enriched binding sites using MACS2 callpeak program ^59^ (Galaxy Version 2.1.1.20160309.6). The resulting bedgraph files were converted to bigwig using ‘Wig/BedGraph-to-bigWig converter’ (Galaxy Version 1.1.1), replicates were combined using ‘Concatenate datasets’ (Galaxy Version 1.0.0). Matrix was computed using ‘computeMatrix’ (Galaxy v.2.3.6.0) in DeepTools ^60^ to prepare data for plotting heat maps and/or profiles of given regions. The genome-wide distributions, heat maps and metagene plots were created using ‘plotHeatmap’ (Galaxy Version 3.1.2.0.1) and ‘plotProfile’ (Galaxy Version 3.1.2.0.0) tools, respectively. The phospho-over-total signal ratios (log_2_-ratio) were calculated using ‘bigwigCompare’ (Galaxy Version 3.1.2.0.0). To generate principal component analysis (PCA) plots ‘plotPCA’ (Galaxy Version 3.1.2.0.0) was used.

*P* values were calculated using Student’s t-test. The ‘‘n’’ values represent number of biological replicates, and the error bars correspond to ± standard deviation (s.d) among biological and technical replicates.

## Data availability

All the raw datasets from sequencing experiments are deposited in NCBI, accession number GSE138548.

## Acknowledgments

We thank N.S. Gray (Dana Farber Cancer Institute) for providing NVP-2, C.J. Hastie (University of Dundee) for anti-phospho-PP4R2-Thr173 antibody, D. Hasson (Mount Sinai) for advice and assistance in analyzing ChIP-seq data, J. Michalak for early efforts to deplete PP1 and N. Jain for technical assistance. This work was supported by National Institutes of Health grant R35 GM127289 to R.P.F. Next-generation sequencing was supported in part by grant P30 CA196521 to the Tisch Cancer Institute.

## Author Contributions

P.K.P designed, performed and analyzed most of the experiments. S.K. performed shRNA and siRNA depletions of PP1 and PP4 subunits and subsequent biochemical analysis. B.B. performed NVP-2 treatment and analysis of Cdk9 target phosphorylation, ChIP-qPCR and initial ChIP-seq analysis of phosphorylated and total Spt5 and Pol II. M.S. generated mammalian PP1 expression constructs and validated PP1 as a Cdk9 substrate. R.P.F. designed and supervised experiments and interpreted data. P.K.P and R.P.F wrote the paper.

## References

1 Buratowski, S. Progression through the RNA polymerase II CTD cycle. Mol Cell 36, 541–546 (2009).

2 Zaborowska, J., Egloff, S. & Murphy, S. The pol II CTD: new twists in the tail. Nat Struct Mol Biol 23, 771–777, doi:10.1038/nsmb.3285 (2016).

3 Buratowski, S. The CTD code. Nat Struct Biol 10, 679–680, doi:10.1038/nsb0903-679 (2003).

4 Egloff, S. & Murphy, S. Cracking the RNA polymerase II CTD code. Trends Genet 24, 280–288, doi:10.1016/j.tig.2008.03.008 (2008).

5 Schwer, B. & Shuman, S. Deciphering the RNA polymerase II CTD code in fission yeast. Mol Cell 43, 311–318 (2011).

6 Hsin, J. P. & Manley, J. L. The RNA polymerase II CTD coordinates transcription and RNA processing. Genes Dev 26, 2119–2137, doi:10.1101/gad.200303.112 (2012).

7 Poss, Z. C. et al. Identification of Mediator Kinase Substrates in Human Cells using Cortistatin A and Quantitative Phosphoproteomics. Cell Rep 15, 436–450, doi:10.1016/j.celrep.2016.03.030 (2016).

8 Sansó, M. & Fisher, R. P. Pause, Play, Repeat: CDKs Push RNAP II's Buttons. Transcription 4(2013).

9 Sansó, M. et al. P-TEFb regulation of transcription termination factor Xrn2 revealed by a chemical genetic screen for Cdk9 substrates. Genes Dev 30, 117–131, doi:10.1101/gad.269589.115 (2016).

10 Adelman, K. & Lis, J. T. Promoter-proximal pausing of RNA polymerase II: emerging roles in metazoans. Nat Rev Genet 13, 720–731 (2012).

11 Core, L. & Adelman, K. Promoter-proximal pausing of RNA polymerase II: a nexus of gene regulation. Genes Dev 33, 960–982, doi:10.1101/gad.325142.119 (2019).

12 Yamaguchi, Y. et al. NELF, a multisubunit complex containing RD, cooperates with DSIF to repress RNA polymerase II elongation. Cell 97, 41–51 (1999).

13 Baluapuri, A. et al. MYC Recruits SPT5 to RNA Polymerase II to Promote Processive Transcription Elongation. Mol Cell, doi:10.1016/j.molcel.2019.02.031 (2019).

14 Glover-Cutter, K. et al. TFIIH-associated Cdk7 kinase functions in phosphorylation of C-terminal domain Ser7 residues, promoter-proximal pausing, and termination by RNA polymerase II. Mol Cell Biol 29, 5455–5464 (2009).

15 Larochelle, S. et al. Cyclin-dependent kinase control of the initiation-to-elongation switch of RNA polymerase II. Nat Struct Mol Biol 19, 1108–1115 (2012).

16 Nilson, K. A. et al. THZ1 Reveals Roles for Cdk7 in Co-transcriptional Capping and Pausing. Mol Cell 59, 576–587, doi:10.1016/j.molcel.2015.06.032 (2015).

17 Peterlin, B. M. & Price, D. H. Controlling the elongation phase of transcription with P-TEFb. Mol Cell 23, 297–305 (2006).

18 Vos, S. M. et al. Structure of activated transcription complex Pol II-DSIF-PAF-SPT6. Nature 560, 607–612, doi:10.1038/s41586-018-0440-4 (2018).

19 Vos, S. M., Farnung, L., Urlaub, H. & Cramer, P. Structure of paused transcription complex Pol II-DSIF-NELF. Nature 560, 601–606, doi:10.1038/s41586-018-0442-2 (2018).

20 Yamada, T. et al. P-TEFb-mediated phosphorylation of hSpt5 C-terminal repeats is critical for processive transcription elongation. Mol Cell 21, 227–237 (2006).

21 Booth, G. T., Parua, P. K., Sanso, M., Fisher, R. P. & Lis, J. T. Cdk9 regulates a promoter-proximal checkpoint to modulate RNA polymerase II elongation rate in fission yeast. Nat Commun 9, 543, doi:10.1038/s41467-018-03006-4 (2018).

22 Jonkers, I., Kwak, H. & Lis, J. T. Genome-wide dynamics of Pol II elongation and its interplay with promoter proximal pausing, chromatin, and exons. Elife 3, e02407, doi:10.7554/eLife.02407 (2014).

23 Glover-Cutter, K., Kim, S., Espinosa, J. & Bentley, D. L. RNA polymerase II pauses and associates with pre-mRNA processing factors at both ends of genes. Nat Struct Mol Biol 15, 71–78 (2008).

24 Davidson, L., Muniz, L. & West, S. 3' end formation of pre-mRNA and phosphorylation of Ser2 on the RNA polymerase II CTD are reciprocally coupled in human cells. Genes Dev 28, 342–356, doi:10.1101/gad.231274.113 (2014).

25 Fong, N., Saldi, T., Sheridan, R. M., Cortazar, M. A. & Bentley, D. L. RNA Pol II Dynamics Modulate Co-transcriptional Chromatin Modification, CTD Phosphorylation, and Transcriptional Direction. Mol Cell, doi:10.1016/j.molcel.2017.04.016 (2017).

26 Proudfoot, N. J. Transcriptional termination in mammals: Stopping the RNA polymerase II juggernaut. Science 352, aad9926, doi:10.1126/science.aad9926 (2016).

27 Parua, P. K. et al. A Cdk9-PP1 switch regulates the elongation-termination transition of RNA polymerase II. Nature 558, 460–464, doi:10.1038/s41586-018-0214-z (2018).

28 Vanoosthuyse, V. et al. CPF-associated phosphatase activity opposes condensin-mediated chromosome condensation. PLoS Genet 10, e1004415, doi:10.1371/journal.pgen.1004415 (2014).

29 Kecman, T. et al. Elongation/Termination Factor Exchange Mediated by PP1 Phosphatase Orchestrates Transcription Termination. Cell Rep 25, 259–269 e255, doi:10.1016/j.celrep.2018.09.007 (2018).

30 Cortazar, M. A. et al. Control of RNA Pol II Speed by PNUTS-PP1 and Spt5 Dephosphorylation Facilitates Termination by a "Sitting Duck Torpedo" Mechanism. Mol Cell 76, 896–908 e894, doi:10.1016/j.molcel.2019.09.031 (2019).

31 Eaton, J. D., Francis, L., Davidson, L. & West, S. A unified allosteric/torpedo mechanism for transcriptional termination on human protein-coding genes. Genes Dev 34, 132–145, doi:10.1101/gad.332833.119 (2020).

32 Pei, Y. & Shuman, S. Characterization of the Schizosaccharomyces pombe Cdk9/Pch1 protein kinase: Spt5 phosphorylation, autophosphorylation, and mutational analysis. J Biol Chem 278, 43346–43356 (2003).

33 Olson, C. M. et al. Pharmacological perturbation of CDK9 using selective CDK9 inhibition or degradation. Nat Chem Biol 14, 163–170, doi:10.1038/nchembio.2538 (2018).

34 Grallert, A. et al. A PP1-PP2A phosphatase relay controls mitotic progression. Nature 517, 94–98, doi:10.1038/nature14019 (2015).

35 Viladevall, L. et al. TFIIH and P-TEFb coordinate transcription with capping enzyme recruitment at specific genes in fission yeast. Mol Cell 33, 738–751 (2009).

36 Ivanov, D., Kwak, Y. T., Guo, J. & Gaynor, R. B. Domains in the SPT5 protein that modulate its transcriptional regulatory properties. Mol Cell Biol 20, 2970–2983 (2000).

37 Larochelle, S. et al. Dichotomous but stringent substrate selection by the dual-function Cdk7 complex revealed by chemical genetics. Nat Struct Mol Biol 13, 55–62 (2006).

38 Bernecky, C., Plitzko, J. M. & Cramer, P. Structure of a transcribing RNA polymerase II-DSIF complex reveals a multidentate DNA-RNA clamp. Nat Struct Mol Biol 24, 809–815, doi:10.1038/nsmb.3465 (2017).

39 Ehara, H. et al. Structure of the complete elongation complex of RNA polymerase II with basal factors. Science 357, 921–924, doi:10.1126/science.aan8552 (2017).

40 Booth, G. T., Wang, I. X., Cheung, V. G. & Lis, J. T. Divergence of a conserved elongation factor and transcription regulation in budding and fission yeast. Genome Res 26, 799–811, doi:10.1101/gr.204578.116 (2016).

41 Gomes, N. P. et al. Gene-specific requirement for P-TEFb activity and RNA polymerase II phosphorylation within the p53 transcriptional program. Genes Dev 20, 601–612 (2006).

42 Vassilev, L. T. et al. In vivo activation of the p53 pathway by small-molecule antagonists of MDM2. Science 303, 844–848, doi:10.1126/science.1092472 (2004).

43 Voss, M. et al. Protein phosphatase 4 is phosphorylated and inactivated by Cdk in response to spindle toxins and interacts with gamma-tubulin. Cell Cycle 12, 2876–2887, doi:10.4161/cc.25919 (2013).

44 Nedea, E. et al. The Glc7 phosphatase subunit of the cleavage and polyadenylation factor is essential for transcription termination on snoRNA genes. Mol Cell 29, 577–587, doi:10.1016/j.molcel.2007.12.031 (2008).

45 Schreieck, A. et al. RNA polymerase II termination involves C-terminal-domain tyrosine dephosphorylation by CPF subunit Glc7. Nat Struct Mol Biol 21, 175–179, doi:10.1038/nsmb.2753 (2014).

46 Grohmann, D. et al. The initiation factor tfe and the elongation factor Spt4/5 compete for the RNAP clamp during transcription initiation and elongation. Mol Cell 43, 263–274 (2011).

47 Baejen, C. et al. Genome-wide Analysis of RNA Polymerase II Termination at Protein-Coding Genes. Mol Cell 66, 38–49 e36, doi:10.1016/j.molcel.2017.02.009 (2017).

48 Shetty, A. et al. Spt5 Plays Vital Roles in the Control of Sense and Antisense Transcription Elongation. Mol Cell 66, 77–88 e75, doi:10.1016/j.molcel.2017.02.023 (2017).

49 Fong, N. et al. Effects of Transcription Elongation Rate and Xrn2 Exonuclease Activity on RNA Polymerase II Termination Suggest Widespread Kinetic Competition. Mol Cell 60, 256–267, doi:10.1016/j.molcel.2015.09.026 (2015).

50 Laitem, C. et al. CDK9 inhibitors define elongation checkpoints at both ends of RNA polymerase II-transcribed genes. Nat Struct Mol Biol 22, 396–403, doi:10.1038/nsmb.3000 (2015).

51 Sen, I. et al. DAF-16/FOXO requires Protein Phosphatase 4 to initiate transcription of stress resistance and longevity promoting genes. Nat Commun 11, 138, doi:10.1038/s41467-019-13931-7 (2020).

52 Blethrow, J. D., Glavy, J. S., Morgan, D. O. & Shokat, K. M. Covalent capture of kinase-specific phosphopeptides reveals Cdk1-cyclin B substrates. Proc Natl Acad Sci U S A 105, 1442–1447 (2008).

53 Yamano, H., Ishii, K. & Yanagida, M. Phosphorylation of dis2 protein phosphatase at the C-terminal cdc2 consensus and its potential role in cell cycle regulation. EMBO J 13, 5310–5318 (1994).

54 Wu, J. Q. et al. PP1-mediated dephosphorylation of phosphoproteins at mitotic exit is controlled by inhibitor-1 and PP1 phosphorylation. Nat Cell Biol 11, 644–651, doi:10.1038/ncb1871 (2009).

55 Swaffer, M. P., Jones, A. W., Flynn, H. R., Snijders, A. P. & Nurse, P. CDK Substrate Phosphorylation and Ordering the Cell Cycle. Cell 167, 1750–1761 e1716, doi:10.1016/j.cell.2016.11.034 (2016).

56 Sansó, M. et al. A Positive Feedback Loop Links Opposing Functions of P-TEFb/Cdk9 and Histone H2B Ubiquitylation to Regulate Transcript Elongation in Fission Yeast. PLoS Genet 8, e1002822 (2012).

57 Mirzoeva, O. K. & Petrini, J. H. DNA replication-dependent nuclear dynamics of the Mre11 complex. Mol Cancer Res 1, 207–218 (2003).

58 Langmead, B. & Salzberg, S. L. Fast gapped-read alignment with Bowtie 2. Nat Methods 9, 357–359, doi:10.1038/nmeth.1923 (2012).

59 Feng, J., Liu, T., Qin, B., Zhang, Y. & Liu, X. S. Identifying ChIP-seq enrichment using MACS. Nat Protoc 7, 1728–1740, doi:10.1038/nprot.2012.101 (2012).

60 Ramirez, F. et al. deepTools2: a next generation web server for deep-sequencing data analysis. Nucleic Acids Res 44, W160–165, doi:10.1093/nar/gkw257 (2016).

