## Supplemental Figures and Table for "Distinct Cdk9-phosphatase switches act at the beginning and end of elongation by RNA polymerase II"

**Running title: Cdk9-phosphatase circuits regulate transcription elongation**

### Supplementary Figures and Legends

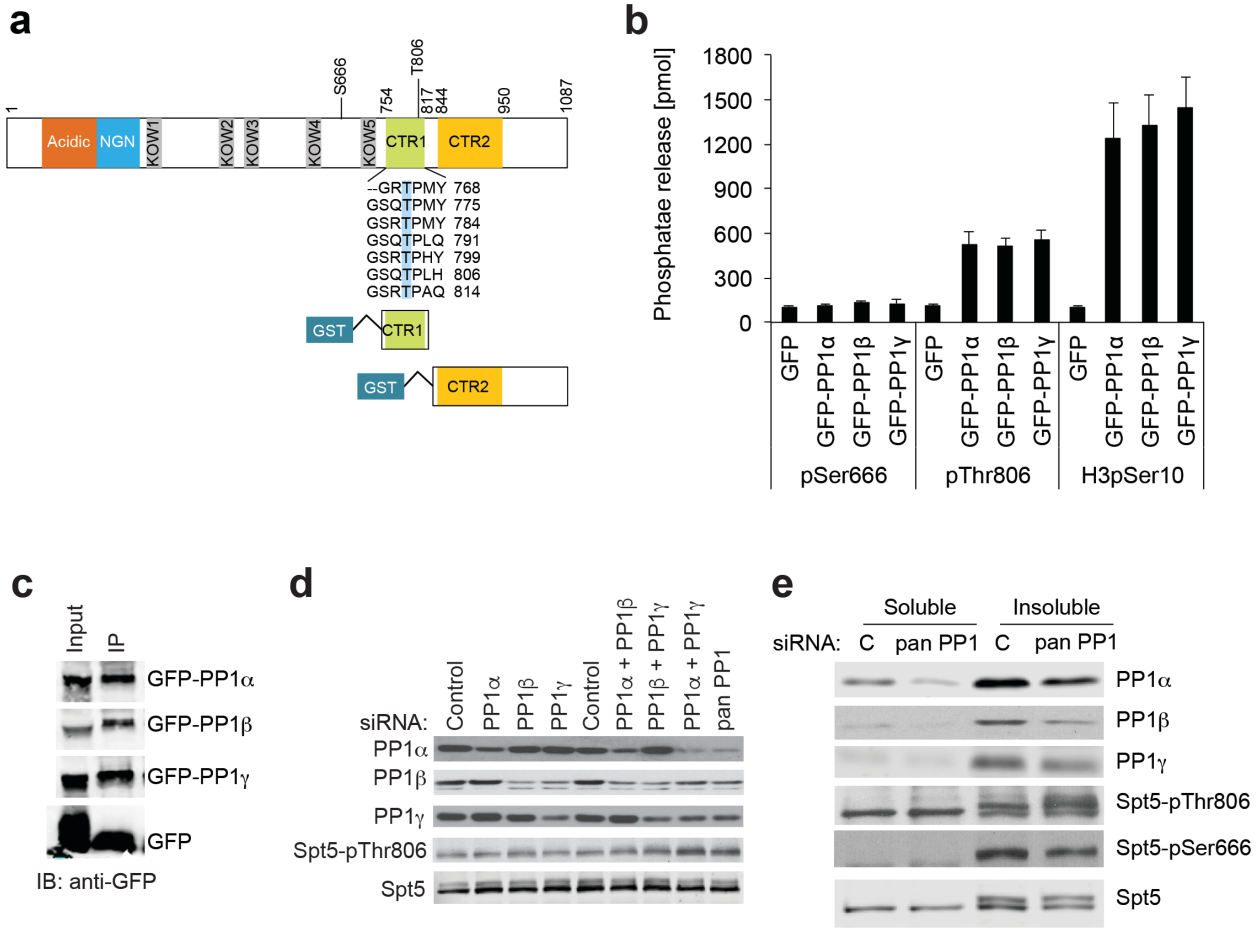


**Supplementary Fig. 1 PP1 regulates Spt5 phosphorylation in human cells. a** Schematic diagram of Spt5 protein. **b** Immunoprecipitated GFP-tagged PP1 isoforms (PP1α, PP1β and PP1γ) dephosphorylated pThr806-containing phosphopeptide and a control (H3pSer10), but not pSer666. Phosphate release was measured colorimetrically. Error bars indicate + s.d. of three biological replicates. **c** Anti-GFP immunoblot shows amounts of PP1 isoforms immunoprecipitated. **d** Immunoblot to measure depletion of PP1 isoforms (PP1α, PP1β and PP1γ) using isoform-specific siRNAs (singly or in combinations) and levels of Spt5, and Spt5-pThr806. **e** Immunoblot of chromatin fraction (insoluble) indicates pThr806 was increased, whereas pSer666 was unchanged in cells depleted of all three PP1 isoforms with an siRNA cocktail.

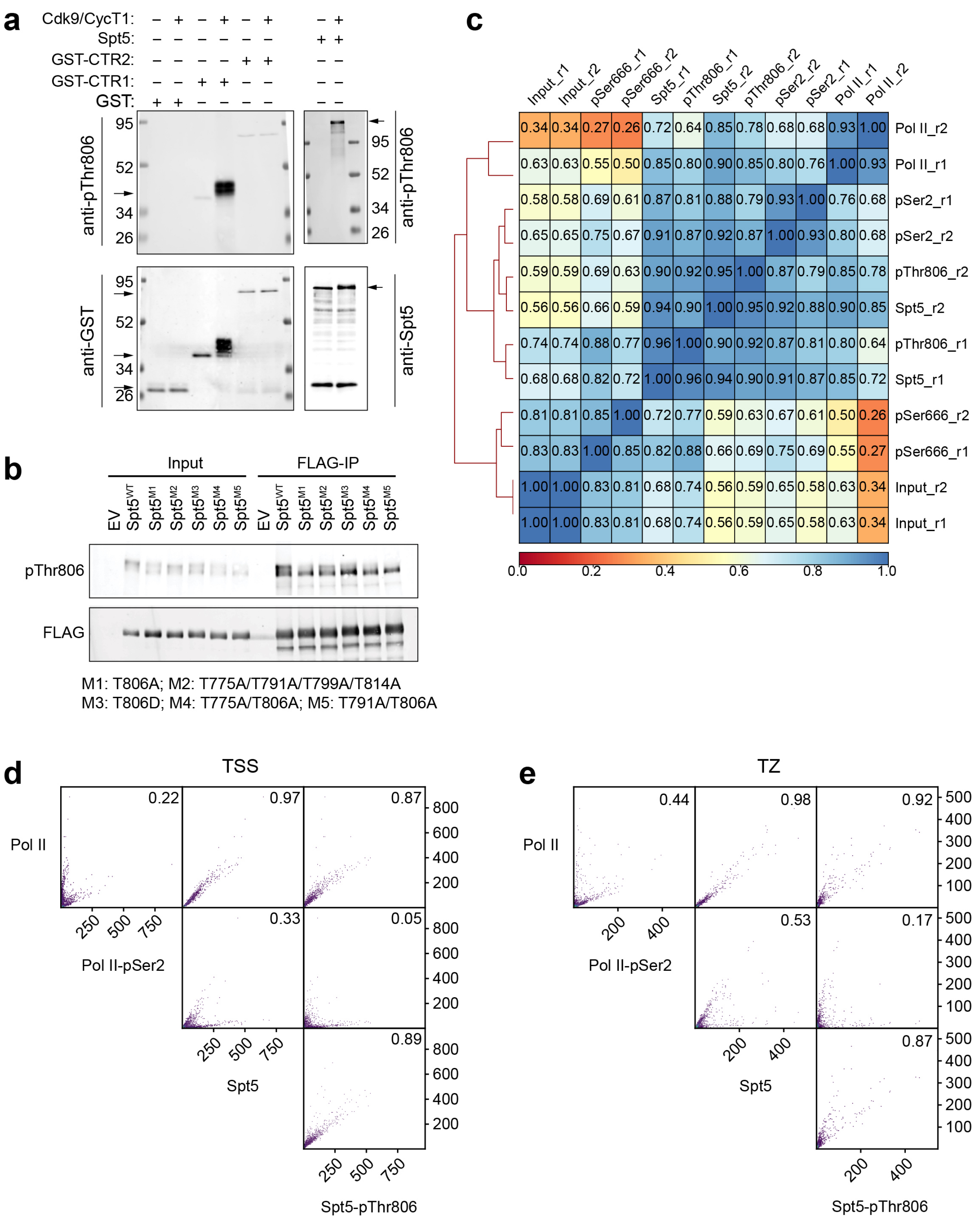


**Supplementary Fig. 2 ChIP-seq analysis of Spt5-pThr806 and Pol II-pSer2. a** Immunoblot analysis shows that upon phosphorylation by Cdk9, anti-pThr806 antibody recognizes GST-CTR1 (amino acids 720-830) and full-length protein, but not GST-CTR2 (amino acids 844-

1087); top (anti-pThr806), bottom left (anti-GST), and bottom right (anti-Spt5). (Note: apparent increase in rabbit anti-GST signal in lane containing GST-CTR1 + Cdk9/cyclin T1 is due to incomplete stripping of membrane after initial probing with rabbit anti-pThr806.) **b** Ectopically expressed FLAG-tagged Spt5—wild-type (WT) and indicated CTR1-mutant variants (M1-M5)—was directly immunoblotted (left) or immunoprecipitated with anti-FLAG (M2) antibody and probed with anti-pThr806 and anti-FLAG antibodies (right). **c** Correlation between ChIP-seq samples. Paired-end sequencing reads were mapped to human genome using Bowtie2. Average read coverages were calculated by feeding resulting BAM files into 'multiBamSummary' program (Galaxy Version 3.1.2.0.0). Values in boxes represent Pearson's correlation coefficients between corresponding samples (*n* = 2 biological replicates). **d** Scatterplots represent correlation among Pol II, Pol II-pSer2, Spt5 and Spt5-pThr806 around the TSS (-50 to +500 bp) for genes highly occupied by Pol II (*n* = 2,572). **e** Scatterplots represent correlation among Pol II, Pol II-pSer2, Spt5 and Spt5-pThr806 at termination zone (TZ; -50 to +5,000 bp of the CPS) for genes highly occupied by Pol II (*n* = 2,572).

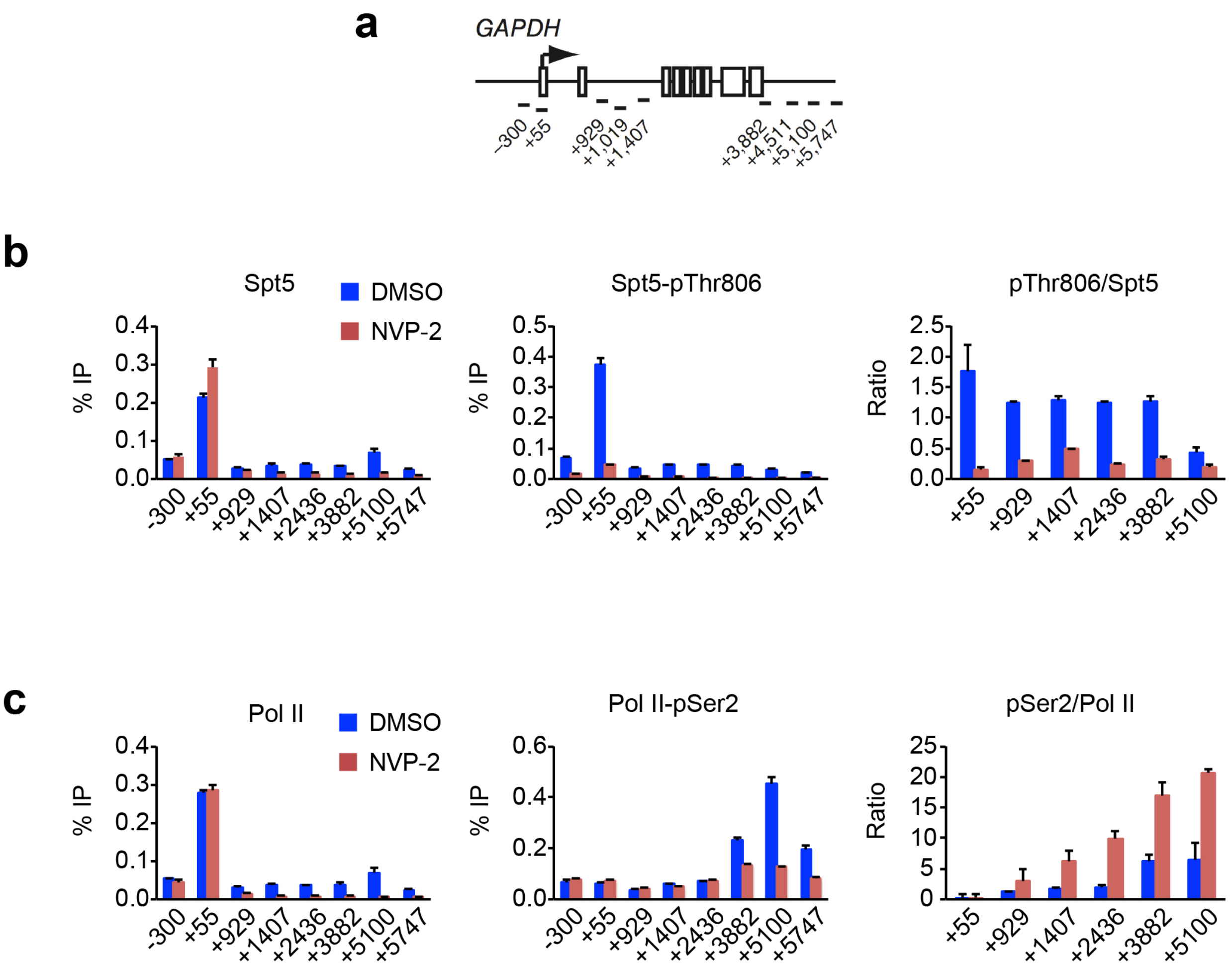


**Supplementary Fig. 3 Cdk9 inhibition diminishes Spt5-pThr806 but increases Pol II CTD-pSer2 on *GAPDH*. a** Schematic of the *GAPDH* gene, indicating positions of primer pairs used in ChIP-qPCR analysis. **b** ChIP-qPCR analysis of total Spt5 and Spt5-pThr806 in HCT116 cells treated with NVP-2 (250 nM) or mock treated (DMSO) for 1 hr. **c** ChIP-qPCR analysis of total Pol II and CTD-pSer2 in HCT116 cells treated with NVP-2 (250 nM) or mock treated (DMSO) for 1 hr. Error bars indicate + s.d. of four biological replicates (**b** and **c**).

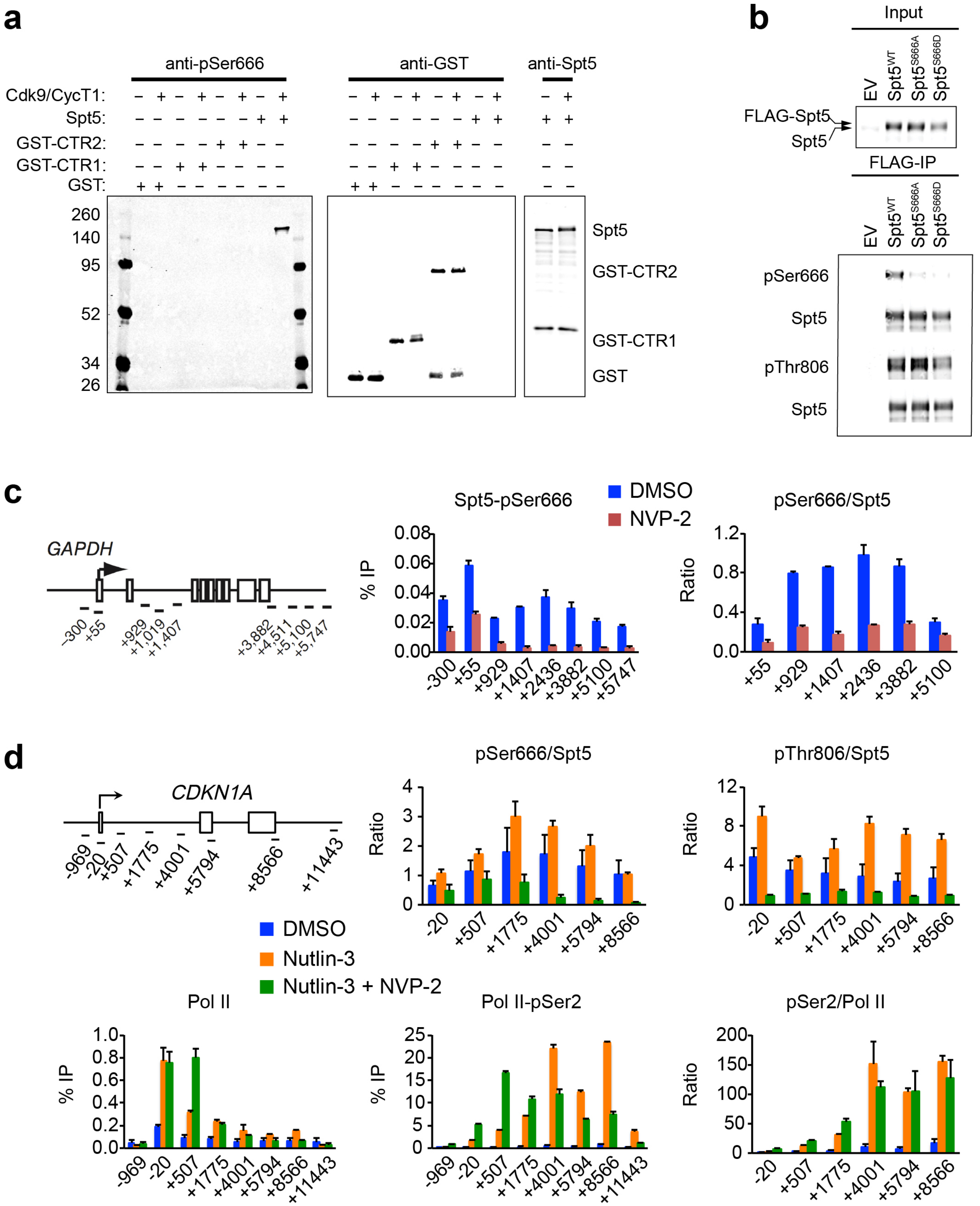


**Supplementary Fig. 4 Distribution and phosphorylation of Spt5 and Pol II-Ser2 on chromatin. a** Immunoblot analysis shows that upon phosphorylation by Cdk9, anti-pSer666 antibody recognizes full-length Spt5 protein, but not GST-CTR1 or GST-CTR2; blots were probed with anti-pSer666 (left), anti-GST (middle), and anti-Spt5 (right). **b** Ectopically expressed FLAG-tagged Spt5—wild-type (WT) and Ser666-mutant variants (Spt5$^{S666A}$ and Spt5$^{S666D}$)—was directly immunoblotted (top, “input”) or immunoprecipitated with anti-FLAG antibody (bottom, “FLAG-IP”) and probed with anti-pSer666, anti-Spt5, anti-pThr806 antibodies. **c** ChIP-qPCR analysis of Spt5-pSer666 on *GAPDH* in HCT116 cells treated with NVP-2 (250 nM) or mock treated (DMSO) for 1 hr. **d** Schematic of *CDKN1A* gene, indicating positions of primer pairs used in ChIP-qPCR analysis, and ChIP-qPCR analysis of Spt5-pSer666, Spt5-pThr806, Pol II and Pol II-pSer2 in HCT116 cells mock treated (DMSO) or treated with nutlin-3 (5 μM) alone or together with NVP-2 (250 nM). (**c** and **d**) Error bars indicate + s.d. from four biological replicates.

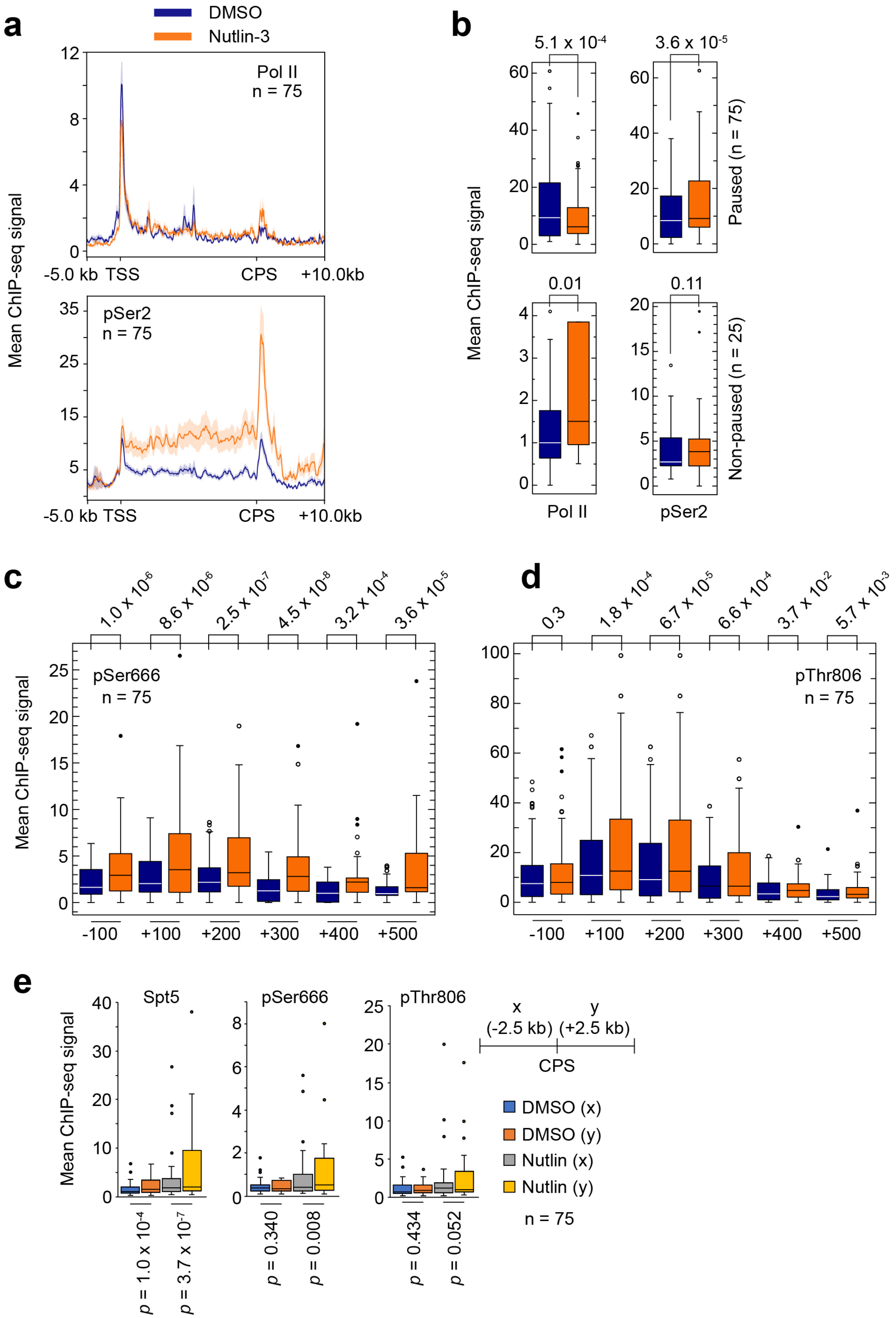

a
DMSO
Nutlin-3
Pol II
n = 75
pSer2
n = 75
Mean ChIP-seq signal
-5.0 kb TSS
CPS
+10.0kb
b
5.1 x 10-4
3.6 x 10-5
0.01
0.11
Paused (n = 75)
Non-paused (n = 25)
Pol II
pSer2
c
1.0 x 10-6
8.6 x 10-6
2.5 x 10-7
4.5 x 10-8
3.2 x 10-4
3.6 x 10-5
pSer666
n = 75
-100
+100
+200
+300
+400
+500
d
0.3
1.8 x 10-4
6.7 x 10-5
6.6 x 10-4
3.7 x 10-2
5.7 x 10-3
pThr806
n = 75
e
Spt5
pSer666
pThr806
p = 1.0 x 10-4
p = 3.7 x 10-7
p = 0.340
p = 0.008
p = 0.434
p = 0.052
x
(-2.5 kb)
y
(+2.5 kb)
CPS
DMSO (x)
DMSO (y)
Nutlin (x)
Nutlin (y)
n = 75

**Supplementary Fig. 5 Distribution of Pol II and Pol II-pSer2 on genes induced by p53 activation. a** Metagene analysis of Pol II and Pol II-pSer2 at genes induced by nutlin-3 ($n$ = 75; paused genes). **b** Box plots comparing ChIP-seq signals in first 100-nt interval downstream of TSS, in the absence or presence of nutlin-3, for Pol II and Pol II-pSer2 at genes induced by nutlin-3, divided into those classified as pause-regulated (Pause index ≥ 2.0, $n$ = 75) or non-paused (pause index < 2.0, $n$ = 25). **c** Box plots show distribution of Spt5-pSer666 ChIP-seq signals in 100-nt intervals from -100 to +500 bp of TSS ($n$ = 75). **d** Box plots show distribution of Spt5-pThr806 ChIP-seq signals in 100-nt intervals from -100 to +500 bp of TSS ($n$ = 75). **e** Box plots show enrichment of Spt5 (left), Spt5-pSer666 (middle) and Spt5-pThr806 (right) mean ChIP-seq signals in intervals around the CPS (x = -2.5 kb to 0; y = 0 to +2.5 kb) when cells were treated with nutlin-3 or mock-treated (DMSO) ($n$ = 75).

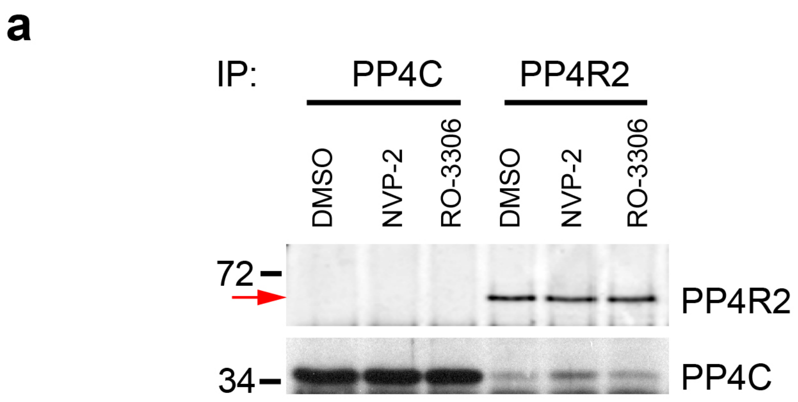


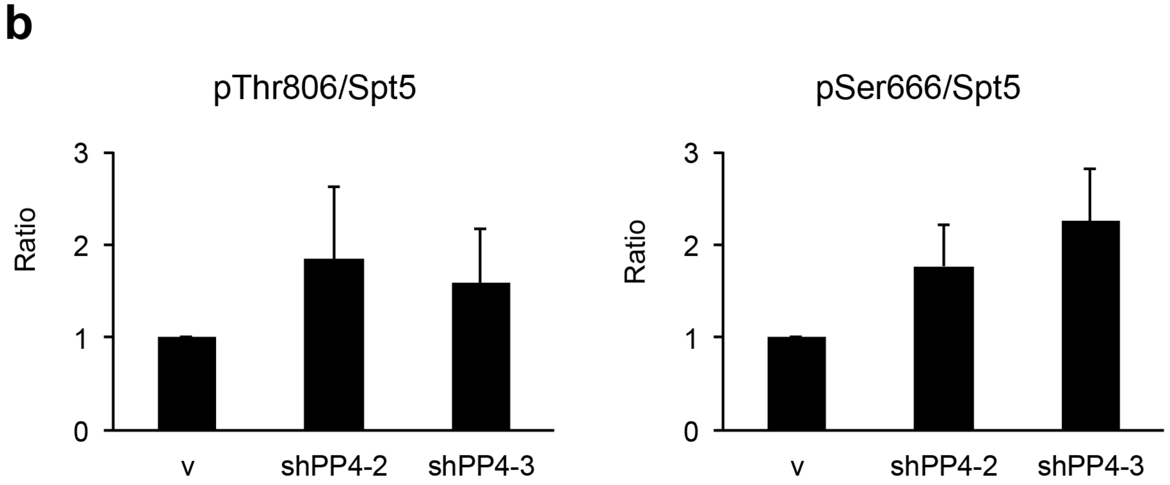


**Supplementary Fig. 6 PP4 is a candidate Spt5 phosphatase. a** PP4C and PP4R2 were immunoprecipitated from extracts prepared from HCT116 cells mock treated (DMSO) or treated with NVP-2 (250 nM) or RO-3306 (10 μM) for 1 hr. PP4C was co-precipitated with anti-PP4R2 but PP4R2 was not detectable in anti-PP4C immunoprecipitate. **b** Quantification of immunoblot results (*n* = 4 independent experiments) in Fig. 6d.

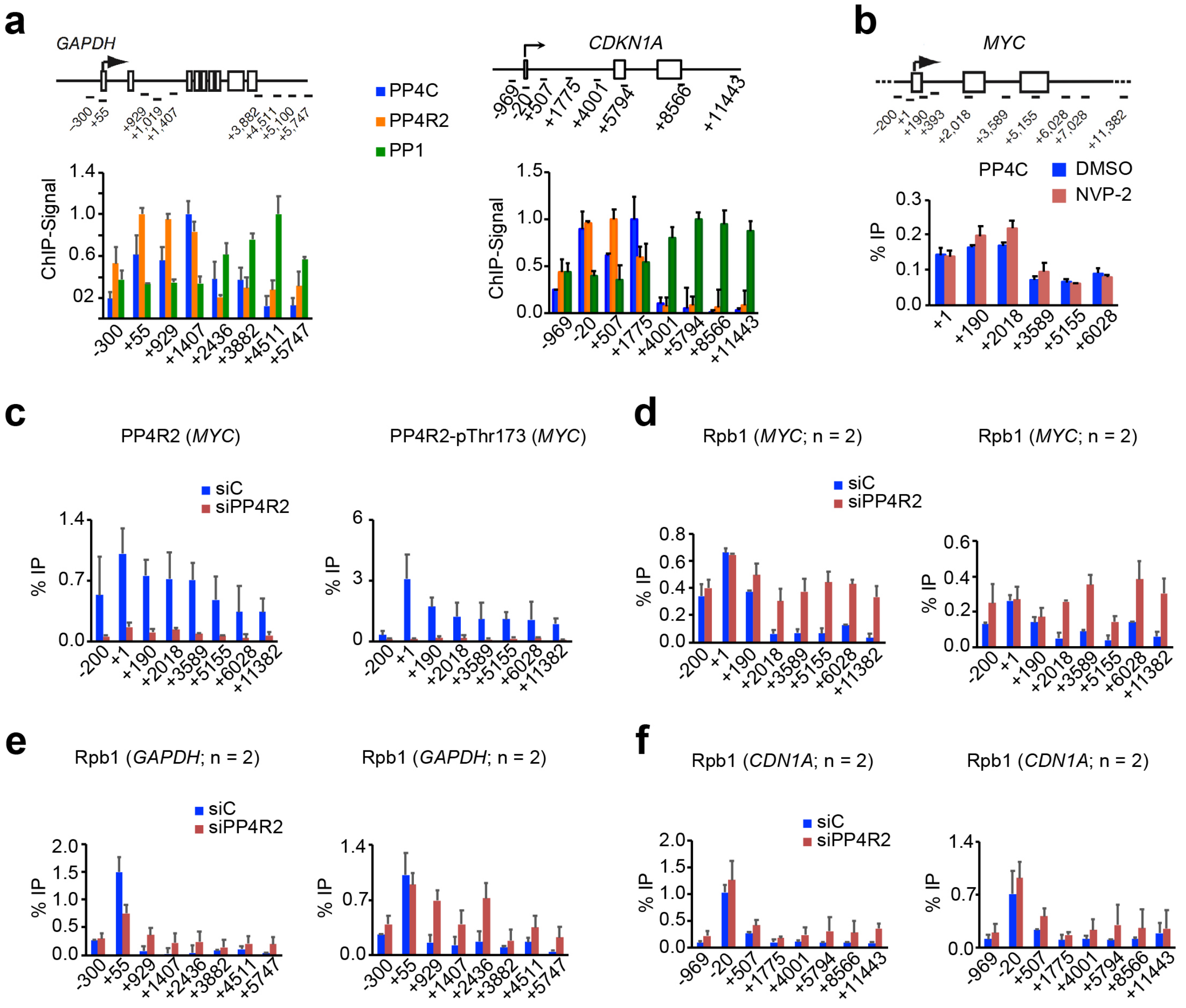


**Supplementary Fig. 7 Distinct spatial distributions and substrate specificities of two Cdk9-regulated, phospho-Spt5 phosphatases. a** Schematic of the *GAPDH* and *CDKN1A* genes, indicating positions of primer pairs used in ChIP-qPCR analysis (*top*). ChIP-qPCR analysis of PP4C, PP4R2 and PP1γ on the *GAPDH* gene in unperturbed HCT116 cells (*bottom*) reveals reciprocal distribution patterns: PP4 subunits are enriched near the 5' end whereas PP1γ occupancy peaks at the 3' end. Error bars indicate + s.d. of two biological replicates. **b** Schematic of the *MYC* gene, indicating positions of primer pairs used in ChIP-qPCR analysis (*top*). Inhibition of Cdk9 with NVP-2 (250 nM, 1 hr) does not affect PP4C occupancy on transcribed chromatin. Error bars indicate + s.d. of two biological replicates. **c** ChIP-qPCR analysis of PP4R2 and PP4R2-pThr173 on the *MYC* gene upon siRNA-mediated depletion of PP4R2 reveals chromatin occupancy of PP4R2 and the Thr173-phosphorylated form is reduced, validating specificity of the anti-pThr173 antibody. Error bars indicate + s.d. of two biological replicates. **d** ChIP-qPCR analysis of Pol II on the *MYC* gene upon siRNA-mediated depletion of PP4R2. Error bars indicate + s.d. of two biological replicates from two separate experiments. **e** ChIP-qPCR analysis of Pol II on the *GAPDH* gene upon siRNA-mediated depletion of PP4R2. Error bars indicate + s.d. of two biological replicates from two separate experiments. **f** ChIP-qPCR analysis of Pol II on the *CDKN1A* gene upon siRNA-mediated depletion of PP4R2. Error bars indicate + s.d. of two biological replicates from two separate experiments.

**Supplementary Table 1. Primers for Site-Directed Mutagenesis**

| Name | Sequence | Key |
|---|---|---|
| SK101 | CTGATCCGGGGAGCCATAGGCGCAAAGCCAC | SUPT5H_S666A_F |
| SK102 | GTGGCTTTGCGCCTATGGCTCCCCGGATCAG | SUPT5H_S666A_R |
| SK103 | GCTGATCCGGGGATCCATAGGCGCAAAGCCACC | SUPT5H_S666D_F |
| SK104 | GGTGGCTTTGCGCCTATGGATCCCCGGATCAGC | SUPT5H_S666D_R |
| PP511 | TCATGCAGGGGCGCCTGTGAGCCGTAG | SUPT5H_T806A-F |
| PP512 | CTACGGCTCACAGGCGCCCCTGCATGA | SUPT5H_T806A-R |
| PP513 | CTGCCATCATGCAGGGGATCCTGTGAGCCGTAGTGTG | SUPT5H_T806D-F |
| PP514 | CACACTACGGCTCACAGGATCCCCTGCATGATGGCAG | SUPT5H_T806D-R |
| PP519 | GCCATACATGGGCGCCTGGGAGCCATACA | SUPT5H_T775A-F |
| PP520 | TGTATGGCTCCCAGGCGCCCATGTATGGC | SUPT5H_T775A-R |
| PP523 | TCCTGGAGGGGTGCCTGTGAGCCGTAC | SUPT5H_T791A-F |
| PP524 | GTACGGCTCACAGGCACCCCTCCAGGA | SUPT5H_T791A-R |
| PP525 | CCGTAGTGTGGGGCGCGGCTACCATCC | SUPT5H_T799A-F |
| PP526 | GGATGGTAGCCGCGCCCCACACTACGG | SUPT5H_T799A-R |
| PP527 | TCTGGGCAGGAGCGCGGCTGCCATC | SUPT5H_T814A-F |
| PP528 | GATGGCAGCCGCGCTCCTGCCCAGA | SUPT5H_T814A-R |
| MS390 | CCTTGGAGGCGCTACAGGTCTCGTGGCATTTG | PPP1CC_ T311A_F |
| MS391 | CAAATGCCACGAGACCTGTAGCGCCTCCAAGG | PPP1CC_T311A_R |
